# Multispecies integrated population model reveals bottom-up dynamics in a seabird predator-prey system

**DOI:** 10.1101/2020.06.26.174250

**Authors:** Maud Quéroué, Christophe Barbraud, Frédéric Barraquand, Daniel Turek, Karine Delord, Nathan Pacoureau, Olivier Gimenez

## Abstract

Assessing the effects of climate and interspecific relationships on communities is challenging because of the complex interplay between species population dynamics, their interactions, and the need to integrate information across several biological levels (individuals – populations – communities). Usually used to quantify species interactions, integrated population models (IPMs) have recently been extended to communities. These models allow fitting multispecies matrix models to data from multiple sources while simultaneously accounting for various sources of uncertainty in each data source. We used multispecies IPMs accommodating climate conditions to quantify the relative contribution of climate vs. interspecific interactions on demographic parameters, such as survival and breeding success, in the dynamics of a predator-prey system. We considered a stage-structured predator–prey system combining 22 years of capture–recapture data and population counts of two seabirds, the Brown Skua (*Catharacta lönnbergi*) and its main prey the Blue Petrel (*Halobaena caerulea*) both breeding on the Kerguelen Islands in the Southern Ocean. Our results showed that climate and predator-prey interactions drive the demography of skuas and petrels in different ways. The breeding success of skuas appeared to be largely driven by the number of petrels and to a lesser extent by intraspecific density-dependence. In contrast, there was no evidence of predation effects on the demographic parameters of petrels, which were affected by oceanographic factors (chlorophyll a and sea surface temperature anomalies). We conclude that bottom-up mechanisms are the main drivers of this skua-petrel system. We discuss the mechanisms by which climate variability and predator-prey relationships may affect the demographic parameters of these seabirds. Taking into account both species interactions and environmental covariates in the same analysis improved our understanding of species dynamics.

## Introduction

The effects of climate changes on the diversity and the structure of communities have been reported repeatedly (Walther et al. 2002, Parmesan 2006, Hoegh-Guldberg and Bruno 2010, Miller et al. 2018). However, the underlying mechanisms remain poorly understood due to the complex dynamics of interacting species: within species, between species and between species and the environment (Godfray and May 2014). Following disturbance, such as changes in environmental conditions, the abundance and distribution of species are expected to be modified according to the position and extent of the species’ niche (Thomas et al. 2004). Because the effects of environmental variability on mortality, fecundity and dispersal may differ between species (Grosbois et al. 2008, Jenouvrier 2013), changes in structure and diversity appear at the community level. However, studying species-by-species responses to environmental changes may overlook the role played by species interactions on those responses, and contribute to a lesser extent to the larger understanding of species interactions that is required by community ecology.

Population dynamics models have been used to understand the effect of interspecific interactions and environment on species demography. However, these models are in general not demographically structured (Stenseth et al. 2015, Pacoureau et al. 2019a, Stoessel et al. 2019) or only partially (Millon et al. 2014, Saunders et al. 2018, Pacoureau et al. 2019b). Unstructured approaches consider individuals as being equivalent but differences in size, age and ontogenic stages exist within a population and may be of importance in the context of interspecific interactions. As argued by Miller and Rudolf (2011), the consideration of the stage structure of populations can lead to a better understanding of community structure and dynamics. Interactions between species such as predation or competition do not necessarily have a homogeneous impact on the different stages of the interacting species. For example, young individuals might be predominantly preyed upon in carnivore–ungulate systems (Gervasi et al. 2015). Therefore, to detect and understand species interactions, we need to consider jointly the demography of several stage-structured populations (Oken and Essington 2015).

Although well developed for single-species dynamics (Tuljapurkar and Caswell 1997, Caswell 2001), demographic stage-structured models have received little attention in community ecology (but see Chu and Adler (2015) on a plant system). The difficulty is that multispecies demography analysis requires integrating information across several biological levels (individual – population – community) which, in turn, requires unifying all available data sources into a single framework. Integrated population models (IPMs) have been recently developed to infer population demography by making complete use of all available sources of information (see Schaub and Abadi 2011, and Zipkin and Saunders 2018 for reviews). In their simplest form, these models combine population counts and demographic data into a single framework, which allows the estimation of demographic parameters while simultaneously accounting for various sources of uncertainty in each data source (e.g. measurement error or parameter estimation) (Besbeas et al. 2002). The IPM framework has been extended to multiple species (Péron and Koons 2012) for competition/parasitism, and more recently for predator-prey interactions (Barraquand and Gimenez 2019).

Here, our main objective was to quantify the relative contribution of environmental changes and species interactions on demographic parameters of a predator and its prey. Therefore, we used a multispecies IPM framework accommodating the effects of local and global climatic conditions on demographic parameters, such as survival and breeding, while explicitly considering species interactions. We applied our approach on a stage-structured predator–prey system combining 22 years of capture-recapture data and population counts on two seabirds, the Brown Skua (*Catharacta lönnbergi*) and its main prey the Blue Petrel (*Halobaena caerulea*) (‘skua’ and ‘petrel’ hereafter) breeding on the Kerguelen Islands in the Southern Ocean.

Because seabirds often occupy high level positions in food-webs, bottom-up forcing which implies population regulation through climate driven limitation in food availability, has long been featured as the dominant paradigm to understand their dynamics (Lack 1967, Aebischer et al. 1990, Stenseth et al. 2002). Seabird foraging behavior and demography reflect the influences of climate variability which directly impacts biological processes in marine ecosystems and cascade through food webs up to seabirds (Barbraud and Weimerskirch 2001, Jenouvrier et al. 2003). However, top-down pressures as predation at breeding colonies are also known to affect the vital rates of seabirds (Hipfner et al. 2012). There is increasing evidence that bottom-up and top-down processes often act in concert and differently affect demographic parameters (Suryan et al. 2006, Horswill et al. 2014, 2016). For example, the effects of predation and resources limitation caused breeding failure of Black-legged Kittiwakes (*Rissa tridactyla*) (Regehr and Montevecchi 1997) and population declines of Arctic Skuas (*Stercorarius parasiticus*) (Perkins et al. 2018). Therefore, quantifying the relative strength of environmental conditions and predator-prey effects is essential for a better understanding of the drivers of population dynamics in seabirds. This is all the more important as climate changes impact the physical properties of the oceans, including the Southern Ocean (Gille 2002, Han et al. 2014) and, through the trophic food web, affect demography and population dynamics of seabirds (Barbraud et al. 2012, Sydeman et al. 2015), including some of the species studied here (Barbraud and Weimerskirch 2003).

Using a multispecies IPM, we assessed the relative contribution of environment and predator-prey interactions on seabirds’ demographic parameters. We estimated survival and adult breeding success for the two interacting species, and assessed the impacts of climatic conditions on these demographic parameters to understand the contribution of predator-prey interactions in shaping population dynamics.

## Materials and Methods

### Study site and Species

Skuas and petrels were studied on Mayes Island (49°28’S, 69°57’E), a 2.8 km² uninhabited island of the Kerguelen Islands in the Southern Ocean where the two species breed during the austral summer (October-February).

The petrel is a small (150–250g) long-lived seabird belonging to the family of *Procellariiformes*. At Kerguelen Islands, petrels feed on macrozooplankton and micronekton feeder, mainly crustaceans and fishes (Cherel et al. 2002, 2014). Individuals from Mayes Island spend the nonbreeding season (from mid-February to September) between the polar front and the northern limit of the pack-ice (57-62°S) between longitudes 20°W and 90°E (Cherel et al. 2016). Birds return to breeding colonies in early September (Quillfeldt et al. 2020). Mayes Island is covered with dry soils and dense vegetation, providing suitable breeding sites for approximately 142,000 breeding pairs of these burrowing petrels (Barbraud and Delord 2006). In late October, a single egg is laid in a burrow dug in peat soil under tall and dense vegetation. The incubation lasts 45-49 days and the chick rearing period 43-60 days (Jouventin et al. 1985). The chick fledges in early February. Both sexes participate in parental care by alternating foraging trips during the incubation and fledging periods.

The skua is a medium sized (1.1 – 2.2 kg) long-lived seabird belonging to the family of *Charadriiformes*. On Mayes Island between 80 and 120 pairs breed annually (Mougeot et al. 1998). Breeding pairs form in October with a high mate fidelity, and generally establish themselves in the same territory each year (Parmelee and Pietz 1987), which they tenaciously defend throughout the breeding season. Generally, two eggs are laid between October and December. The incubation lasts 28-32 days and the chicks rearing period 40-50 days (Higgins and Davied 1996). Skuas are extremely plastic in their foraging techniques and adapt their diet depending on the local availability of prey (Carneiro et al. 2015). On Mayes Island, during the breeding season, Blue Petrels represent 95% of the skua diet (Pacoureau et al. 2019c). Skuas from Mayes Island overwinter in the southern hemisphere between 10°E and 150°E (Delord et al. 2018).

During the breeding period on Mayes Island, the predation of petrels by skuas takes place mainly at night, when petrels come out or arrive at their burrows (Mougeot and Bretagnolle 2000a). Skuas mostly prey on petrels on the ground, but they can also catch petrels in flight (Mougeot et al. 1998, Pacoureau et al. 2019c). Vocalizing petrels, especially those without partners, are more easily detected by skuas during the courtship period (Mougeot and Bretagnolle 2000b). Skuas may also prey on chicks during the fledging period.

### Count and capture-recapture data

Data of both skuas and petrels were collected during the breeding seasons from 1996/1997 to 2017/2018. For convenience, breeding seasons are named from 1996 to 2017 hereafter. The time interval used in our model starts before the wintering of species and ends at the end of the breeding period. Two types of data were used: count data corresponding to the number of burrows or territories occupied by seabirds and capture-recapture (CR) data of adult seabirds found on the monitored area. Each year, adult individuals of both species were checked at specific times following the species phenology to determine the breeding status of each bird. The breeding status of marked birds was determined at the end of the breeding period. In the following we describe how the data were collected for the two species. For clarity, all parameters for skuas are indicated by *S* and by *P* for petrels.

Around 200 individually marked burrows of petrels were inspected each year from early-to-mid November just after the egg-laying to check for eggs and to identify marked adults, and then in late January just before fledging of the chicks. Each year since 1985 (see Barbraud and Weimerskirch 2005), all fledglings as well as new individuals found in burrows were marked with a stainless steel band (captured by hand, marked, and replaced in their burrow). Petrels never observed with an egg or a chick during a given breeding season were considered as nonbreeders (*NB*). Individuals were identified as breeders if they laid a single egg or raised a chick and as successful breeders if their chick fledged (SB). Two categories of failed breeders were used depending on the stage of failure: egg stage (*FBE*) or chick stage (*FBC*). Given that the first sampling period occurred just after laying, it is very unlikely that nonbreeders were failed breeders. These breeding statuses allowed the construction of the individual capture histories (*Ch*_*p*_ and constituted our CR data. The annual number of adult petrels (*Y*_*p*_), *i.e.* count data, was estimated as the number of occupied burrows. Each occupied burrow was considered as being frequented by a pair of petrels. We considered that this count included all adult individuals, both breeders and non-breeders.

For skuas, each year since 1991, the eastern side of Mayes Island was inspected to identify territories of skuas. A territory was considered established when a pair strongly defended an area against other skuas (Mougeot et al. 1998). Around 50 nesting territories were visited four to eight times from mid-October (after egg-laying) to late-February (just before fledging of the chicks) each year. Chicks just before fledging, as well as new adult individuals, were marked with a metal ring and a plastic ring to facilitate individual identification using binoculars. Breeding status was determined by checking the nest contents for the presence of eggs or young chicks. Skuas never observed with an egg or a chick were considered as nonbreeders (*NB*). Individuals were identified as breeders if they laid at least one egg or raised a chick. If the eggs did not hatch or the chicks died, both members of the pair were considered as failed breeders (*FB*). Given that the first sampling period occurred just after laying, it is very unlikely that nonbreeders represented failed breeders. Successful breeders were defined as individuals that fledged one or two chicks, and were denoted as *SB*1 or *SB*2, respectively. These breeding statuses allowed the construction of the individual capture histories (*Ch*_*S*_) and constituted our CR data. The annual number of skuas (*Y*_*S*_), *i.e.* count data, was estimated as the number of territories and each territory was considered occupied by a pair of skuas. We considered that this count included all adult individuals, both breeders and non-breeders.

For both species, individual breeding status could be considered as “uncertain” (*C*) in case of difficulties to assign their breeding status (lack of information, missed checks, individuals never re-observed). Only adult individuals that have bred at least once between the 1996 and the 2017 breeding seasons were kept in the data set for analysis to eliminate potential transient individuals (*n* = 318 for skuas and *n* = 1210 for petrels). Individual capture histories (*Ch*) started at their first breeding attempt recorded. Based on the high probability of observing breeders in the study site, we assumed that the first breeding attempt was correctly detected. New individuals found in monitored burrows or territories are considered as immigrants to the study site (*N*_*im*_).

The presence of chicks was used to assign a breeding status to adult individuals captured in the breeding area. In order to maintain the independence of the data, we did not include information on chicks in the model. Therefore, the fecundity was a fixed value. We considered one chick for each pair of seabird, considered as successful breeders (*N*_*SB,P*_) for petrels or successful breeders with one chick (*N*_*SB*1,*S*_) for skuas (*f*_*SB,P*_ and *f*_*SB*1,*S*_ are equal to 1, respectively). For skuas that successfully fledged two chicks (*f*_*SB*2,*S*_ we considered two chicks per pair of skuas (*f*_*SB*2,*P*_ is equal to 2). Since juveniles only return to the breeding sites as adults to attempt to breed for the first time (from four year old or older), we did not have data on juvenile states.

### Integrated Population Model

We built a two-species IPM that combines count and CR data and allows estimating abundances and demographic rates (Besbeas et al. 2002, Schaub and Abadi 2011). More specifically, we connected two IPMs, one for predatory skuas and one for petrels, their main prey, through explicit predator-prey relationships (Barraquand and Gimenez 2019). We incorporated the effects of predation within species-specific vital rates such as survival and breeding parameters. This IPM is structured by states which represent life history states (Fig. 1). We built two likelihoods, one for the CR data and the other for the count data which we combined into a joint likelihood.

**Figure 1:**
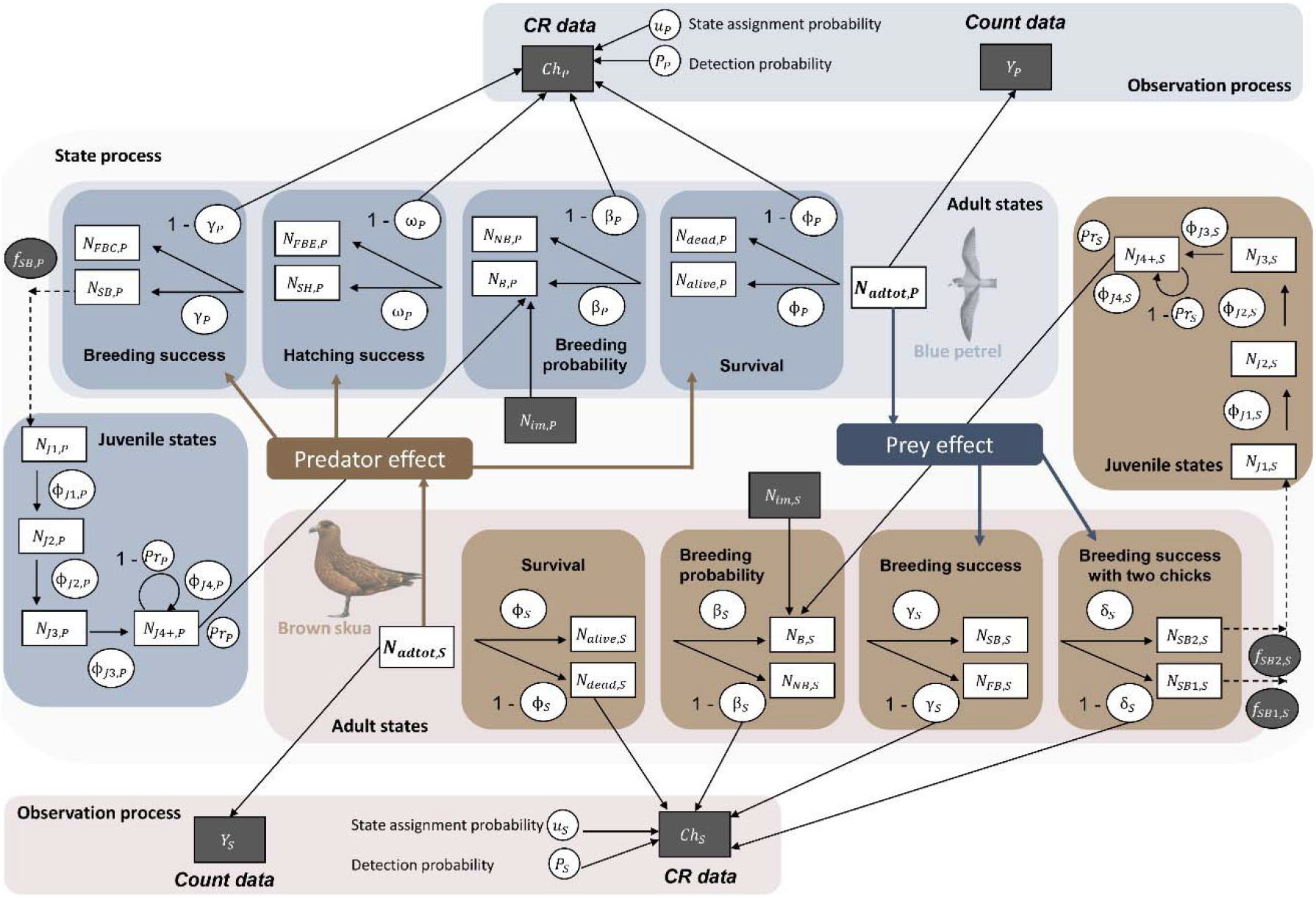
Structure of the multispecies Integrated Population Model. Squares represent the state variables, circles represent the parameters. Data and fixed values are represented with a dark background, estimated state variables and parameters with a white background. Two types of data are used, capture histories (*Ch*) from capture-recapture data and count data (*Y*). Adult apparent survival (*ϕ*), breeding probability (*β*), hatching success (*ω*), breeding success (*γ*), breeding success with two chicks (*δ*), juvenile apparent survival for one to four years old and older (*ϕ*_*J*1_ to *ϕ*_*J*4_), probability of first reproduction (*Pr*), state assignment probability (*u*) and detection probability (*p*) are parameters estimated in the model. Fecundity (*f*) is fixed. The number of adults (*N*_*adtot*_), dead (*N*_*dead*_), alive (*N*_*alive*_), breeders (*N*_*B*_), nonbreeders (*N*_*NB*_), failed breeders (*N*_*FB*_), failed breeders at the stage egg (*N*_*FBE*_), breeders with an egg hatched (*N*_*SH*_), failed breeders at the stage chick (*N*_*FBC*_), successful breeders (*N*_*SB*_), successful breeders with one chick (*N*_*SB*1_) or with two chicks (*N*_*SB*2_) and the number of juveniles of one year old to four years old and older (*N*_*J*1_ to *N*_*J4+*_) are state variables estimated by the model. The number of immigrants (*N*_*im*_) is a fixed vector. The blue part is for Blue Petrels and the brown part is for Brown Skuas. Interspecific relationships are represented with thick arrows.

In the following, we detail the state process following a biological timeline and we explain the different likelihood used. The structure was the same for the two species but states differed in relation to species biology (Fig. 1). The two main differences were: (1) skuas could have up to two chicks versus only one for petrels, (2) the failed-breeder stage in petrels could be split further according to the timing of failure (failure at the incubation vs. chick-rearing stage). For clarity, parameters are indexed by *S* (for skuas) or *P* (for petrels) when differences occur, or by *X* (for *S* or *P*) when the structure is the same for both species. We used Poisson (*Po*) and binomial (*Bin*) distributions to account for demographic stochasticity. Notations of all parameters and state variables are detailed in Appendix S1: Table S1.

### State process

#### Offspring production

The estimated number of skuas and petrels in their first year *i.e.* between 0 and 1 year old (*N*_*J*1,*S*,*t*_ at year *t*, is modelled with a Poisson distribution:

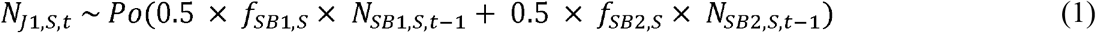

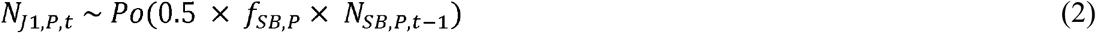

with *N*_*J*1,*S*_ the number of chicks produced by all successful skua breeders (*N*_*SB*1,*S*_ and *N*_*SB*2,*S*_) according to their fecundity (*f*_*SB*1,*S*_: 1 chick and *f*_*SB*2,*S*_: 2 chicks per female skua, sex ratio: 0.5). For petrels, *N*_*J*1,*P*_, is also Poisson distributed but with only one chick (*f*_*SB,P*_ per estimated successful female breeder (*N*_*SB,P*_ with a sex ratio of 0.5).

#### Juvenile survival

The number of juveniles between one and two years *N*_*J*2_, two and three years *N*_*J*3_, and three and four years *N*_*J*4_, are modelled with binomial distributions:

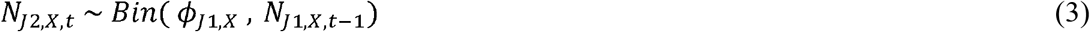

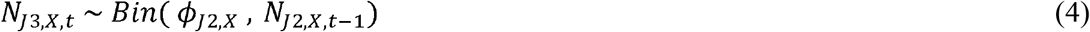

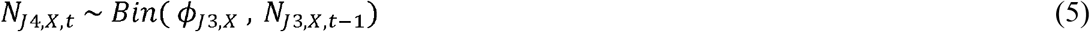

with the apparent survival between one and two years (*ϕ*_*J*1_), between two and three years (*ϕ*_*J*2_ and between three and four years (*ϕ*_*J*3_ respectively. As we observed only adult breeding birds, we had no information on the juvenile phase. We assumed that juvenile apparent survival increased with age (Greig et al. 1983, Grande et al. 2009, Fay et al. 2015), as experienced birds are on average more effective in foraging (Daunt et al. 2007), in competing with conspecifics or in avoiding predators:

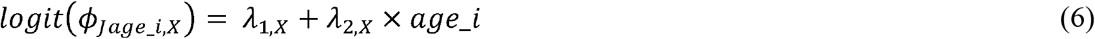

where (*ϕ*_*J*_ is the juvenile apparent survival, age_*i* the age of the juvenile state (from (*N*_*J*1_ to *N*_*J*4_), *λ*_1_ the intercept and *λ*_2_ the slope which is constrained to be positive.

#### Juvenile first breeding attempt

The first breeding attempt in skuas and petrels could start from age four. Four years old individuals and older that did not attempt to breed are in the state *N*_*J*4+_. The individuals that attempted to breed for the first time with a first breeding attempt probability *Pr* are in the state *N*_*J4B*_ and the individuals that did not attempt to breed are in the state *N*_*J4NB*_ :

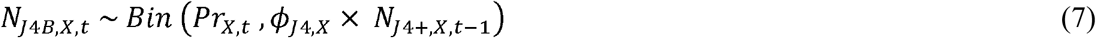

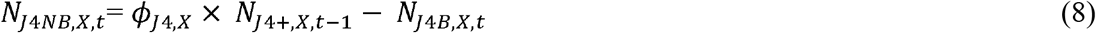

with *ϕ*_*J*4_ the apparent survival for the *N*_*J4+*_ state. The *N*_*J4+*_ state includes individuals that did not attempt to breed (*N*_*J4NB*_) and individuals aged between three and four years *N*_*J*4_ :

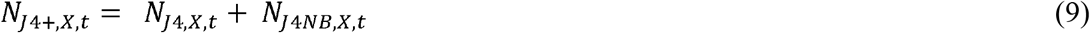

#### Adult survival

For the two species, we modelled the number of surviving adults *N*_*alive*_ at year *t* among the total number of adult individuals (*N*_*adtot*_ at year *t-1* with a binomial distribution, with *ϕ* the adult apparent survival:

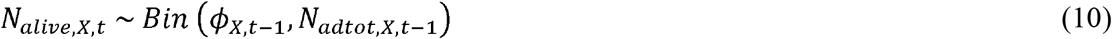

#### Breeding probability

The number of adult individuals that have bred or not bred among those that survived *N*_*alive*_ is modelled as:

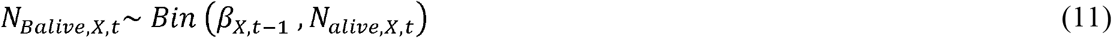

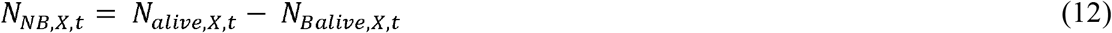

with *β* the probability of breeding, *N*_*Balive*_ the number of adult breeders that survived and *N*_*NB*_ the number of adult nonbreeders. As capture histories started at their first breeding attempt recorded, the immigrants, *i.e.* newly marked individuals (*N*_*B*_) coming for the first time in the colony, were considered as breeders. Then, the total number of breeders (*N*_*B*_) corresponds to the sum of the number of adult breeders that survived (*N*_*Balive*_, the number of immigrants (*N*_*im*_) and the number of juveniles attempting to breed for the first time *N*_*J4B*_ :

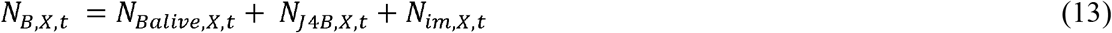

The total number of adults (*N*_*adtot*_) corresponds to the sum of nonbreeders (*N*_*NB*_) and breeders *N*_*B*_ :

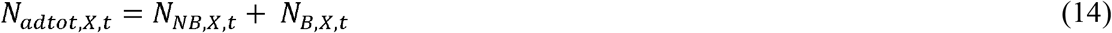

#### Breeding success

Breeding success and failure are modelled differently for skuas and petrels. For skuas, the numbers of failed breeders (*N*_*FB,S*_) and successful breeders (*N*_*SB,S*_ are modelled following a binomial distribution:

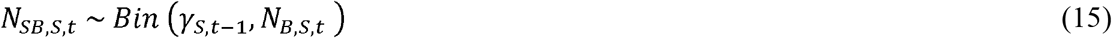

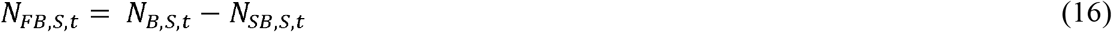

with *γ*_*S*_ the probability of a successful breeding. A successful breeder can then have one or two chicks, respectively *N*_*SBl,S*_ and *N*_*SB*2,*S*_ and this is modelled following a binomial distribution:

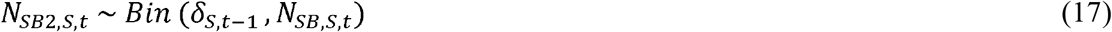

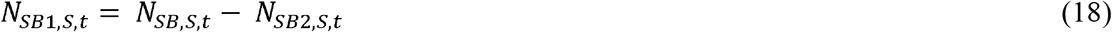

with *δ*_*S*_ the probability of producing two chicks rather than one among the successful breeders.

For petrels, there are two states for failed breeders: one with petrels that failed to hatch their egg (named failed breeder at the egg stage *N*_*FBE,P*_) and the second with petrels that failed to fledge their chick (named failed breeder at the chick stage *N*_*FBC,P*_). Hence, there is a parameter of successful hatching (*δ*_*P*_). The numbers of petrels with an egg that successfully hatched (*N*_*SH,P*_), and the failed breeders at the egg stage (*N*_*FBE,P*_) were modelled following a binomial distribution:

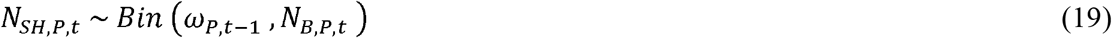

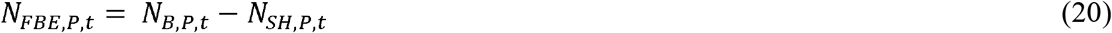

with the probability of successful hatching. Successful breeders (*N*_*SB,P*_) and failed breeders at the chick stage (*N*_*FBC,P*_) were modelled following a binomial distribution:

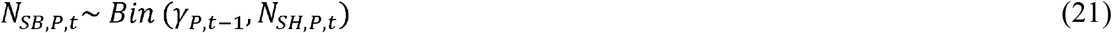

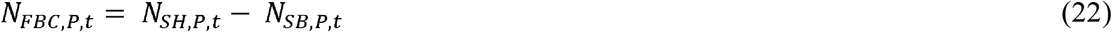

with *y*_*P*_ the probability of successful breeding.

### Count data

The observation equation links the observed adult population count (*Y*) (*i.e.* the number of territories/burrows multiplied by two for a pair of seabird) with the true adult population siz (*N*_*adtot*_), with an additional term for observation error:

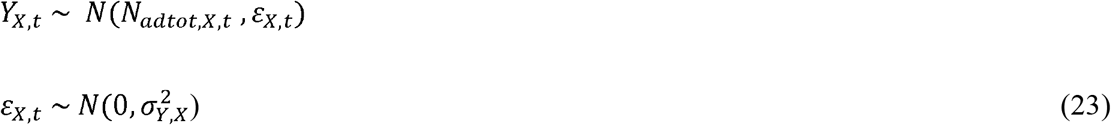

where ∊ is the error term and 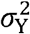 its variance. As only the adult states were observed on the field, we excluded the juvenile states from the observation equation. The likelihood for the population count data is denoted as *L*_*CO,S*_(*Y*_*S*_|*ϕ*_*J*1,*S*_,*ϕ*_*J2,S*_, *ϕ*_*J3,S*_, *ϕ*_*J4,S*_, *Pr*_*S*_, *ϕ*_*S*_, *β*_*S*_, *γ*_*S*_, *δ*_*S*_, *N*_*adtot,S*_) for skuas and as *L*_*CO,P*_(*Y*_*P*_|*ϕ*_*J*1,*P*_, *ϕ*_*J2,P*_, *ϕ*_*J3,P*_, *ϕ*_*J4,P*_, *Pr*_*P*_, *ϕ*_*P*_, *β*_*P*_, *ω*_*P*_, *γ*_*P*_, *N*_*adtot*_,*P*) for petrels.

### Capture-recapture data

For adult CR data, we used multievent capture–recapture models to estimate the demographic parameters (Pradel 2005). These models take into account the imperfect detectability of the individuals as well as the uncertainty in the assignment of states to individuals (Gimenez et al. 2012).

For skuas, our multievent model includes five states: NB, FB, SB1, SB2, dead, and six events: not seen, seen as NB, seen as FB, seen as SB1, seen as SB2, seen as C. For petrels, the five states are: NB, FBE, FBC, SB, dead, and the six events are: not seen, seen as NB, seen as FBE, seen as FBC, seen as SB, seen as C. The following demographic parameters were estimated for the two species: the adult apparent survival probability (*ϕ*_*X*_), the breeding probability (*β*_*X*_), the probability of successful breeding (*γ*_*X*_). The probability of successful breeding with two chicks (*δ*_*S*_) was also estimated for skuas, as well as the probability of hatching (*ω*_*X*_) for petrels. Two additional parameters were also estimated: the detection probability (*P*_*X*_) and the state assignment probability of individuals with uncertain state (*u*_*X*_) . All parameters were time-varying through a yearly random effect, except *u* (Table 1). State transitions were set to be state dependent according to the breeding status in the previous breeding season (Table 1): Breeder 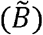 representing birds that attempted to breed the previous breeding season (*FB*, *SB*1, *SB*2 for skuas or *FBE*, *FBC*, ss for petrels) or Nonbreeder 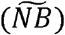 representing birds that already bred previously but did not attempt to breed during the previous breeding season (*NB*). The detection probability and the state assignment probability also depended on the breeding status (Table 1). The likelihood for the CR data for skuas is denoted as *L*_*Cr,S*_(*Ch*_*S*_|*ϕ*_*S*_,*β*_*S*_, *γ*_*S*_, *δ*_*S*_, *P*_*S*_, *u*_*S*_) and *L*_*Cr,P*_(*Ch*_*P*_|*ϕ*_*P*_,*β*_*P*_, *γ*_*P*_, *δ*_*P*_, *P*_*P*_, *u*_*P*_) for petrels.

**Table 1:** Summary of the demographic parameters and their specificities (year random effect or state dependence) for the two species: the Brown Skua (top) and the Blue Petrel (bottom). Notations are 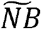: Nonbreeder the previous year, 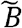: Breeder the previous year, NB: Nonbreeder, FB: Failed Breeder, SB1: Successful Breeder with one fledged chick, SB2: Successful Breeder with two fledged chicks, FBE: Failed Breeder at the Egg stage, FBC: Failed Breeder at the Chick stage and SB: Successful Breeder.

| Species | Parameter | Year random effect | State dependence |
| --- | --- | --- | --- |
| **Skua** | Adult apparent survival $\phi_S$ | ? | $\widetilde{NB}_S \vert \widetilde{B}_S$ |
| | Breeding probability $\beta_S$ | ? | $\widetilde{NB}_S \vert \widetilde{B}_S$ |
| | Breeding success $\gamma_S$ | ? | $\widetilde{NB}_S \vert \widetilde{B}_S$ |
| | Breeding success 2 chicks $\delta_S$ | ? | $\widetilde{NB}_S \vert \widetilde{B}_S$ |
| | Detection probability $p_S$ | ? | $\widetilde{NB}_S \vert \widetilde{B}_S$ |
| | Uncertain state assignment probability $u_S$ | ? | $NB_S \vert FB_S \vert SB1_S \vert SB2_S$ |
| **Petrel** | Adult apparent survival $\phi_P$ | ? | $\widetilde{NB}_P \vert \widetilde{B}_P$ |
| | Breeding probability $\beta_P$ | ? | $\widetilde{NB}_P \vert \widetilde{B}_P$ |
| | Hatching success $\omega_P$ | ? | $\widetilde{NB}_P \vert \widetilde{B}_P$ |
| | Breeding success $\gamma_P$ | ? | $\widetilde{NB}_P \vert \widetilde{B}_P$ |
| | Detection probability $p_P$ | ? | $\widetilde{NB}_P \vert \widetilde{B}_P$ |
| | Uncertain state assignment probability $u_P$ | ? | $NB_P \vert FBE_P \vert FBC_P \vert SB_P$ |

### Joint likelihood

The joint likelihood of the skua IPM is the product of the likelihood for the count data (*L*_*co,S*_) and CR data *L*_*cr,S*_) :

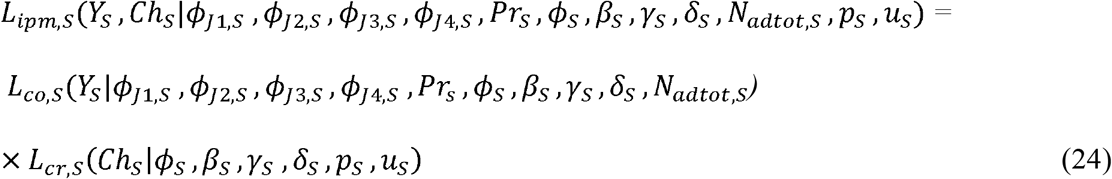

For petrels, the product of the likelihood for the count data *L*_*co,P*_), and CR data *L*_*cr,S*_) is denoted as: *L*_*ipm,P*_(*Y*_*P*_,*Ch*_*P*_|*ϕ*_*Jl,P*_ *ϕ*_*J2,P*_ *ϕ*_*J3,P*_ *ϕ*_*J4,P*_ *Pr*_*P*_, *ϕ*_*P*_, *β*_*P*_*ω*_*P*_, *γ*_*P*_, *N*_*adtot,P*_ *P*_*P*_, *u*_*P*_).

### Interspecific relationships, intraspecific density-dependence, and environmental covariates

We used different covariates to investigate their effects on the adult demographic parameters estimated for the two species (Table 2). We focused only on the demographic parameters of adult individuals because only adults were observed on the field. We tested interspecific predator-prey relationships between skua and petrel, and intraspecific relationships with density-dependence for both species. Moreover, we considered several climatic covariates that were suspected to affect demographic parameters of skuas and petrels, the Southern Annular Mode (SAM) on a large scale, and the Sea Surface Temperature anomalies (SSTa) and Chlorophyll a concentration (Chla) on a local scale. In the following, we provide more details on covariates and how they may affect the demography of skuas and petrels.

**Table 2:**
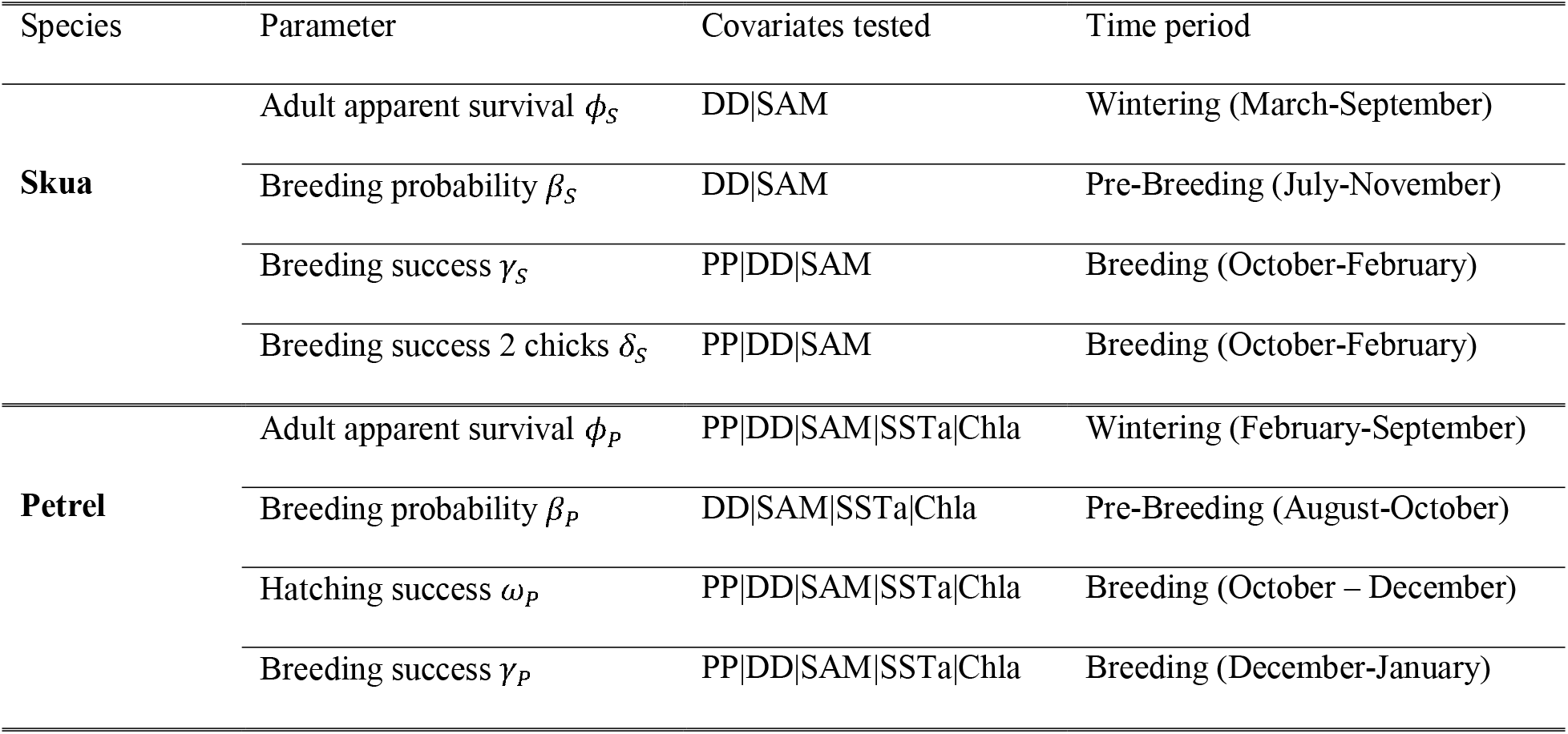
Summary of the covariates tested on the demographic parameters of the two species – the Brown Skua (top) and the Blue Petrel (bottom) – and the time period (in months) considered for each demographic parameter. Notations are PP: Predator-Prey interactions, DD: intraspecific Density-Dependence, SAM: Southern Annual Mode, SSTa: Sea Surface Temperature anomalies, Chla: Chlorophyll a concentration.

| Species | Parameter | Covariates tested | Time period |
| --- | --- | --- | --- |
| **Skua** | Adult apparent survival $\phi_S$ | DD\|SAM | Wintering (March-September) |
| | Breeding probability $\beta_S$ | DD\|SAM | Pre-Breeding (July-November) |
| | Breeding success $\gamma_S$ | PP\|DD\|SAM | Breeding (October-February) |
| | Breeding success 2 chicks $\delta_S$ | PP\|DD\|SAM | Breeding (October-February) |
| **Petrel** | Adult apparent survival $\phi_P$ | PP\|DD\|SAM\|SSTa\|Chla | Wintering (February-September) |
| | Breeding probability $\beta_P$ | DD\|SAM\|SSTa\|Chla | Pre-Breeding (August-October) |
| | Hatching success $\omega_P$ | PP\|DD\|SAM\|SSTa\|Chla | Breeding (October – December) |
| | Breeding success $\gamma_P$ | PP\|DD\|SAM\|SSTa\|Chla | Breeding (December-January) |

### Predator-prey interactions

Multispecies IPMs allow us to explicitly include interspecific relationships between vital rates of one species and estimated population sizes of the other. Based on the high proportion of petrels in the diet of the skuas during the breeding season (Mougeot et al. 1998, Pacoureau et al. 2019c), we predicted that petrel adult apparent survival (*ϕ*_*P*_) should decrease with the number of skuas. As skuas prey on adults and chicks during the fledging period, we predicted that the hatching success (*ω*_*P*_) and fledging success (*γ*_*P*_) would be impacted by the number of predators.

Inversely, we predicted that a large number of petrels in the breeding colony would provide enough food resources for skua and then be favorable to their breeding success (*γ*_*P*_) and breeding success with two chicks (*δ*_*S*_).

### Intraspecific density-dependence

We investigated the effect of intraspecific density-dependence on the demography of the two species as higher density of individuals on the breeding area can lead to and increasing competition for food resources or for territories. Skuas are highly territorial and defend their territories vigorously during the whole breeding season. The most violent fights may even lead to their death. Moreover, the limited number of territories could cause emigration of skuas without territory. Thus, we predicted that the apparent survival (*ϕ*_*S*_), *i.e.* the joint estimation of the mortality and emigration, would be negatively impacted by the number of skuas. This limited number of territories could also lead to a negative density-dependence relationship between breeding probability (*β*_*S*_) and population density. The energetic cost and the time spent in defending a territory throughout the breeding season may limit the time spent searching for food, potentially limiting energy investment in reproduction. We thus predicted a negative effect of population density on the successful breeding parameter (*γ*_*S*_) and the probability to have two chicks rather than one for successful breeders (*δ*_*S*_). For petrels, we also tested the effects of intraspecific competition for food resources, which could affect their adult apparent survival (*ϕ*_*P*_) and their breeding parameters: breeding probability (*β*_*P*_), hatching (*ω*_*P*_) and fledging success (*γ*_*P*_).

### Environmental covariates

Climate variability impacts biological processes in marine ecosystems, which cascade through food webs and are integrated by seabirds (Barbraud and Weimerskirch 2001, Jenouvrier et al. 2003). Hence, we considered several covariates that are suspected to affect populations of petrels and skuas through these bottom-up mechanisms. All covariates are used as proxies of food availability at sea at different scales. In the following, we explain how environmental conditions may impact the two species based on their diet and distribution.

Because skuas have broad wintering areas (Delord et al. 2018), we tested a large-scale environmental covariate, the SAM. In contrast with their diet during the breeding season specialized on the Blue Petrel, during winter skuas adopt a mixed diet composed of low trophic level preys, such as macrozooplankton and crustaceans (Delord et al. 2018). We hypothesized that availability of food resources at sea during the austral winter might have an effect on the body condition of skuas and then affect the survival of skuas. Moreover, skuas may experience a carry-over effect as the additional energy invested by individuals to maintain themselves during poor wintering conditions may have repercussion on their ability to breed the next breeding season (Harrison et al. 2011, Bogdanova et al. 2017).

For petrels, the wintering areas have been determined (Cherel et al. 2016) allowing us to test two covariates used at local scale, the SSTa and the Chla, in addition of the SAM. As their diet is mainly composed of crustaceans and fish feeding at low trophic levels (Cherel et al. 2002, 2014), the food availability at sea may impact the survival of petrels. Moreover, during the breeding season, male and female petrels take turns, one incubating the egg and fasting and the other foraging at sea, which results in substantial variation in their body mass (Chaurand and Weimerskirch 1994a, 1994b, Weimerskirch et al. 1994, Chastel et al. 1995). Therefore, high food availability at sea may allow a good foraging success of the foraging partner that may return to land after a short stay at sea, allowing a good synchronization of the breeding partners on the nest. In contrast, poor conditions could increase the time spent at sea by the foraging partner, which would increase desertion of the nest by the fasting partner and then, reduce the breeding success. We thus predicted that conditions at sea during the breeding season would also affect the breeding success of petrels.

#### Southern Annular Mode

The SAM is a large-scale climate index. SAM is the leading mode of climate variability over the Southern Hemisphere. SAM is defined as the difference of atmospheric pressure between the 40°S and 65°S latitudes (Marshall 2003). SAM influences surface wind, sea surface temperature (SST) and surface chlorophyll concentration. A large majority of the skuas from Mayes Island overwinter north of the polar front (Delord et al. 2018). In the subtropical zone, SAM positive phases induced warm SSTa, low surface chlorophyll concentration and easterly winds driving Ekman transport (the 90° wind-driven net transport on the sea surface), while in the Subantarctic zone there is a convergence of waters that increase downwelling and positive SSTa (Lovenduski and Gruber 2005). We thus predicted that the positive phases of SAM, potentially leading to poorer food availability in the areas used by skuas during the nonbreeding period, would have negative impacts on skua survival and limit their ability to breed the next breeding season. South of the polar front, where petrels spend the winter, positive phases of the SAM are associated with westerly winds. This induces cold SSTa, increased equatorward Ekman transport and drives increased upwelling (Lovenduski and Gruber 2005). Consequently, the biological productivity and potential prey availability for petrels are higher during positive phases of the SAM. We thus predicted that the positive phases of SAM would be favorable for petrel demographic parameters. Data were obtained from the online database of the British Antarctic Survey (http://www.nerc-bas.ac.uk/icd/gjma/sam.html).

#### Sea Surface Temperature anomalies

SSTa reflect local oceanographic conditions that influence the whole marine trophic food web. High SST generally reduces vertical mixing and provides poor growing conditions for zooplankton communities which, through bottom-up mechanisms, induces reduced trophic resources for seabirds (Barbraud et al. 2012, Sydeman et al. 2015). Consequently, year-to-year variation of SST was previously found to be negatively correlated with petrel body condition (Guinet et al. 1998). Therefore, we predicted that high SSTa would negatively affect overwinter survival and breeding success of petrels. The SSTa data were downloaded from the National Oceanic and Atmospheric Administration (“data: NOAA NCEP EMC CMB GLOBAL Reyn_SmithOIv2 monthly ssta”) from 1996 to 2018.

#### Chlorophyll a

Chlorophyll a lies at the bottom of the marine food web and provides resources for higher trophic organisms up to seabirds. Because petrel diet is mainly composed of crustaceans and fish feeding at low trophic levels (Cherel et al. 2002, 2014), we predicted that high concentrations of Chla would be favorable to the survival and breeding success of petrels. The Chla data were downloaded from the NASA Ocean Data with a 9km mapped concentration data of chlorophyll a for the years 1997 to 2001 and from the Nasa Earth Observation (NEO AQUA/MODIS data) monthly for the years 2002 to 2018.

### Assessing the effect of environmental covariates and population densities

We fitted a single multispecies IPM including all the biologically relevant effects. Logit-linear regressions were used to estimate the effect of environmental (SAM, SSTa and Chla) and inter- and intra-specific interactions on demographic parameters (adult apparent survival, breeding probability, hatching probability, breeding success) (Table 2). We used state variables *N*_*adtot,S*_ and *N*_*adtot*_,*P*, respectively the number of adult skuas and petrels, to assess the effects of inter- and intra-specific interactions. For example, we modelled the hatching probability for petrels that bred the previous year 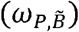, using a logit link:

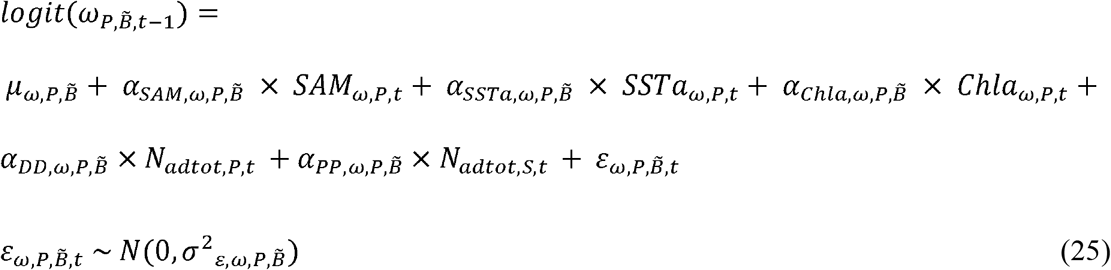

with 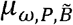 the intercept, 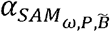 the slope for the climatic covariate 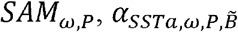 the slope for the climatic covariate 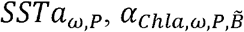 the slope for the climatic covariate 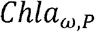 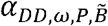 the slope indicating the strength of the intraspecific density-dependence with *N*_*adtot*_,*P* the number of adult petrels 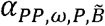 the slope indicating the strength of the predator-prey relationship with *N*_*adtot,S*_ the number of adult skuas, 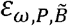 is a yearly random effect and a 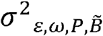 its temporal variance. The descriptions of all logit-linear relationships used on demographic parameters are available in Appendix S2.

For local covariates (SSTa and Chla), we calculated the average values of the covariates in the areas in which petrels were located (Cherel et al. 2016) in a specific time period during which the environment might affect the demographic parameter under investigation (Table 2). Each environmental covariate was standardized to have zero mean and unit variance. However, the inter- and intra-specific covariates were not standardized prior to the analyses because the population sizes were estimated step by step each year. To compare the relative contribution of the effects of each covariate, we calculated the standardized effect of population size (for inter- and intra-specific relationship) posterior to the analyses by multiplying their slopes by the standard deviation of the estimated population sizes. Then, we compared the relative contribution of each covariate using the regression estimate which we used as a measure of effect size.

We computed the 95% and 80% credible intervals (CRI) for the regression coefficients α. We did not interpret uncertain effects (*i.e.* 80% CRI including zero) and focused particularly on clear effects whose sign could be reliably assessed (*i.e.* 95% CRI excluding zero).

### Model implementation

To fit the juvenile apparent survival parameters increasing with age, we modelled them as a positive linear function of age by assigning to the slope *λ*_2_ a *U*(0,1) prior, and by defining the intercept *λ*_1_ with a normal *N*(0,1) prior. The probability of the first breeding attempt (*Pr*) is time-dependent with a uniform prior: *Pr*_*t*_~U(0,1). Normal priors *N*(0,10^4^) were asigned to the regression coefficients (*α*) of the covariate effects. For the variance of the random year effects 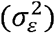 we used a *U*(0,10) vague prior. The state assignment probability of individuals with uncertain state parameter (*u*) was defined *a priori* with a *U*(0,10) vague prior.

Bayesian posterior distributions were approximated *via* Markov chain Monte Carlo (MCMC) algorithms. Two independent MCMC chains of 200,000 iterations were used with a burn-in period of 100,000. One out of five iterations was kept and final inferences were derived from a sample of 2 × 20,000 iterations that resulted from merging the two chains. Gelman-Rubin convergence diagnostic (Brooks and Gelman 1998) was below 1.5 for each parameter and the mixing of the chains was satisfactory. We performed the analyses using Nimble (de Valpine et al. 2017; version 0.9.1) and program R (R Core Team 2020; R version 4.0.3). Code and data are available on GitHub at https://github.com/maudqueroue/MultispeciesIPM_SkuaPetrel.

## Results

### Predator-prey relationships

We estimated positive relationships between two breeding parameters of skuas and the number of adult petrels. The breeding success for at least one chick 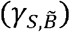 [slope mean 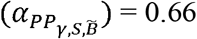 95% CRI (0.37, 1.05)] (Fig. 2a) and the breeding success with two chicks 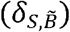 [slope mean 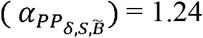; 95% CRI (0.59, 2.12)] (Fig. 2b) for skuas that were breeders the previous breeding season increased with an increasing number of prey. Even though the effects were less clear (95% CRI including zero), the breeding success of petrels tended to be positively impacted by the number of predators (Table 3). We detected a positive relationship between the number of adult skuas and the hatching success of petrels that were breeders the previous breeding season 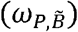, and with the breeding success of petrels that were nonbreeders the previous breeding season 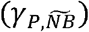. We found no other interspecific relationship on the other parameters (Table 3).

**Table 3:** Regression coefficients estimates for the relationships between covariates (DD: intraspecific Density-Dependence, PP: Predator-Prey interactions, SAM: Southern Annular Mode, SSTa: Sea Surface Temperature anomalies, Chla: Chlorophyll a concentration) and demographic parameters (*ϕ*: adult apparent survival, *β*: breeding probability, *γ*: breeding success, *δ*: breeding success with two chicks, w: hatching success) for Brown Skuas (top) and Blue Petrels (bottom), 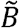: breeders or 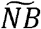: non breeders the previous years. 80% credible intervals that do not include zero are in bold.

| SKUA | DD |  |  |  | PP |  |  |  | SAM |  |  |  |
| --- | --- | --- | --- | --- | --- | --- | --- | --- | --- | --- | --- | --- |
| Parameters | slope | sd | 10% | 90% | slope | sd | 10% | 90% | slope | sd | 10% | 90% |
| $\phi_{S,\tilde{B}}$ | -0.09 | 0.11 | -0.22 | 0.06 | | | | | -0.38 | 0.41 | -0.88 | 0.12 |
| $\phi_{S,\tilde{NB}}$ | -0.12 | 0.15 | -0.28 | 0.09 | | | | | **1.25** | **0.96** | **0.14** | **2.42** |
| $\beta_{S,\tilde{B}}$ | 0.23 | 0.22 | -0.08 | 0.51 | | | | | 0.73 | 0.69 | -0.13 | 1.61 |
| $\beta_{S,\tilde{NB}}$ | 0.11 | 0.19 | -0.16 | 0.33 | | | | | 0.80 | 0.65 | 0.00 | 1.60 |
| $\gamma_{S,\tilde{B}}$ | **-0.39** | **0.14** | **-0.56** | **-0.22** | **0.66** | **0.17** | **0.45** | **0.89** | 0.12 | 0.20 | -0.14 | 0.37 |
| $\gamma_{S,\tilde{NB}}$ | -0.23 | 0.28 | -0.54 | 0.15 | 0.20 | 0.39 | -0.26 | 0.65 | 0.21 | 0.54 | -0.45 | 0.86 |
| $\delta_{S,\tilde{B}}$ | **-0.53** | **0.24** | **-0.85** | **-0.22** | **1.24** | **0.38** | **0.78** | **1.74** | 0.02 | 0.41 | -0.49 | 0.53 |
| $\delta_{S,\tilde{NB}}$ | -0.40 | 0.44 | -0.95 | 0.16 | 0.71 | 0.96 | -0.47 | 1.80 | -1.30 | 2.33 | -4.07 | 0.79 |

| PETREL | DD |  |  |  | PP |  |  |  | SAM |  |  |  | SSTa |  |  |  | Chla |  |  |  |
| --- | --- | --- | --- | --- | --- | --- | --- | --- | --- | --- | --- | --- | --- | --- | --- | --- | --- | --- | --- | --- |
| Parameters | slope | sd | 10% | 90% | slope | sd | 10% | 90% | slope | sd | 10% | 90% | slope | sd | 10% | 90% | slope | sd | 10% | 90% |
| $\phi_{P,\tilde{B}}$ | -0.14 | 0.30 | -0.51 | 0.24 | 0.25 | 0.24 | -0.06 | 0.56 | -0.34 | 0.39 | -0.83 | 0.16 | -0.22 | 0.24 | -0.53 | 0.07 | -0.51 | 0.47 | -1.08 | 0.05 |
| $\phi_{P,\tilde{NB}}$ | -0.38 | 1.05 | -1.67 | 1.04 | 0.43 | 0.49 | -0.25 | 1.01 | 1.27 | 2.26 | -1.11 | 4.51 | 0.01 | 0.96 | -1.01 | 1.25 | -0.56 | 1.70 | -2.43 | 1.59 |
| $\beta_{P,\tilde{B}}$ | **0.51** | **0.25** | **0.19** | **0.86** | | | | | -0.34 | 0.39 | -0.85 | 0.14 | 0.07 | 0.22 | -0.22 | 0.34 | 0.34 | 0.38 | -0.16 | 0.82 |
| $\beta_{P,\tilde{NB}}$ | 0.21 | 0.33 | -0.22 | 0.67 | | | | | 0.40 | 0.38 | -0.07 | 0.85 | **0.36** | **0.22** | **0.09** | **0.63** | **0.63** | **0.40** | **0.12** | **1.13** |
| $\omega_{P,\tilde{B}}$ | 0.12 | 0.25 | -0.18 | 0.43 | **0.28** | **0.14** | **0.11** | **0.44** | -0.10 | 0.25 | -0.41 | 0.20 | 0.08 | 0.16 | -0.12 | 0.28 | -0.35 | 0.29 | -0.70 | 0.01 |
| $\omega_{P,\tilde{NB}}$ | -0.47 | 0.52 | -1.08 | 0.08 | 0.26 | 0.33 | -0.16 | 0.66 | -0.17 | 0.61 | -0.87 | 0.53 | -0.12 | 0.47 | -0.56 | 0.37 | -0.51 | 0.69 | -1.30 | 0.28 |
| $\gamma_{P,\tilde{B}}$ | 0.05 | 0.61 | -0.68 | 0.81 | -0.12 | 0.31 | -0.49 | 0.27 | 0.32 | 0.46 | -0.22 | 0.93 | -0.44 | 0.42 | -0.98 | 0.05 | -0.13 | 0.64 | -0.91 | 0.70 |
| $\gamma_{P,\tilde{NB}}$ | **-0.79** | **0.59** | **-1.53** | **-0.12** | **0.46** | **0.33** | **0.01** | **0.89** | -0.43 | 0.45 | -0.99 | 0.08 | -0.34 | 0.51 | -0.99 | 0.25 | -0.85 | 0.79 | -1.84 | 0.06 |

**Figure 2:**
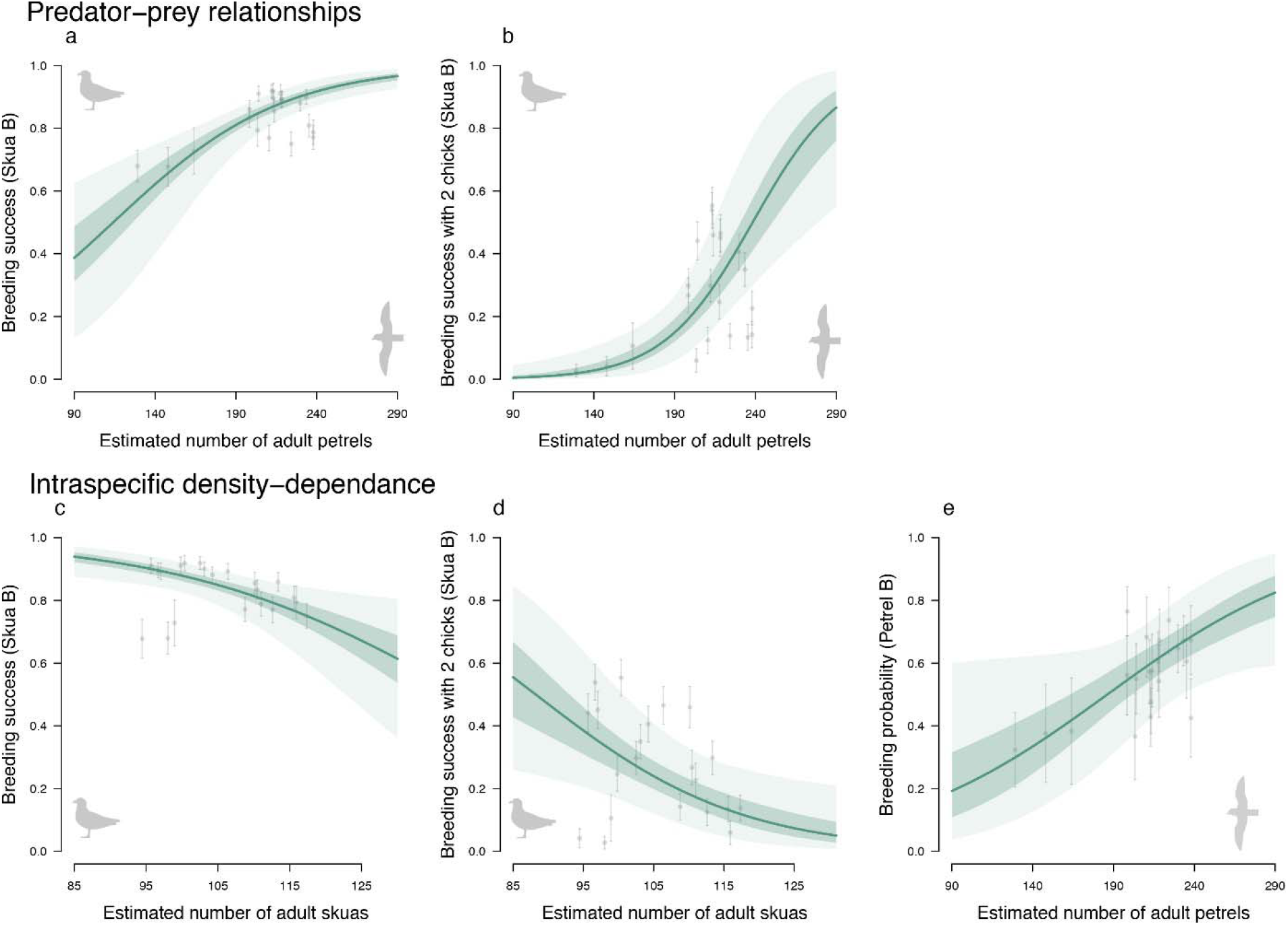
Effects of predator-prey relationships (top panels) and intraspecific density-dependence (bottom panel) on adult demographic parameters for the two seabirds, the Brown Skua and the Blue Petrel. Solid lines represent the estimated relationship between the covariates and the demographic parameters. Shaded areas are the 50% and 95% credibility intervals. Points represent demographic parameter estimates each year (21 years) plotted against covariate. Error bars are standard deviation. Prey effect on (a) the estimated breeding success probability 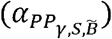 and (b) breeding success with two chicks for skuas that bred the previous breeding season 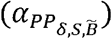. Intraspecific density-dependence effect on (c) the breeding success 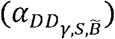 and on (d) breeding success of skuas that were breeders the previous breeding season 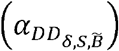 and (e) on the breeding probability of petrels that bred the previous breeding season 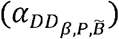.

### Intraspecific density-dependence

The number of skuas had a clear effect on two demographic parameters, namely the breeding success and the breeding success with two chicks for skuas that were breeders the previous breeding season. We found negative density-dependence for the breeding success 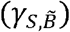 [slope mean 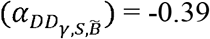; 95% CRI (−0.65, −0.12)] (Fig. 2c) and for the probability of producing two chicks rather than one 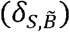 [slope mean 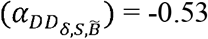; 95% CRI (−1.02, −0.10)] (Fig. 2d). These two breeding parameters were also affected by interspecific relationships and we observed that the predator-prey effects were stronger than intraspecific effects 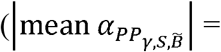 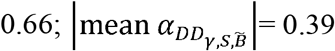 respectively) for the breeding success and 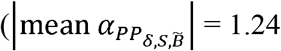 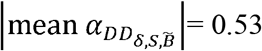 respectively) for the breeding success with two chicks (Table 3).

For petrels, we estimated a positive effect of increased number of adult petrels on the breeding probability for individuals that were breeders the previous breeding season 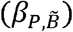 [slope mean 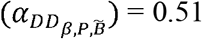; 95% CRI (0.03, 0.98)] (Fig. 2e). Moreover, the number of petrels tended to negatively affect the breeding success of petrels that did not bred the previous breeding season 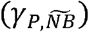 (Table 3).

### Environmental covariates

We found ecologically relevant relationships between environmental covariates and demographic parameters of the two species (Table 3). For petrels, we found positive relationships between the two local environmental covariates (SSTa and Chla) and the breeding probability for individuals that were nonbreeders the previous breeding season 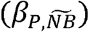. The effect of these environmental covariates on the breeding probability was stronger for the Chla covariate than for the SSTa covariate (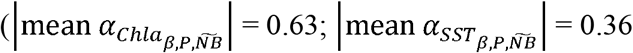 respectively). We estimated a positive relationship between the SAM covariate and the apparent survival of skuas that were nonbreeders the previous breeding season 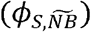.

In addition to the results above, we also estimated the demographic parameters and the number of individuals in each state for both species from 1996 to 2017 (see Appendix S3: Figs. S1– S6).

## Discussion

In this paper, we provide the first application of a multispecies IPM in a predator-prey context. Joint analysis of empirical data for two seabird species allowed us to estimate demographic parameters and population size for both simultaneously. The key advantage of using a multispecies IPM was that it enabled us to use the population sizes estimated by the model for one species to analyze its effect on the demographic parameters of the other species while propagating all sources of uncertainty. Hence, it allowed us to understand the contribution of interspecific interactions on the demographic parameters while further taking into account the effects of climatic conditions. Our results showed that the demography of the predatory skua was mainly driven by the number of petrel prey during the breeding season and by the environment during the nonbreeding season, whereas petrels were mostly impacted by the environment. This suggests that this predator-prey system is mainly driven by bottom-up processes and density-dependent processes.

### Effects of predator-prey relationships

The number of prey is a determining factor in the breeding success of skuas according to our results. Food availability is known to be positively related with breeding parameters in seabirds (Cairns 1988, Piatt et al. 2007, Oro et al. 2014). As diet of skuas during the breeding period is dominated by petrels (Mougeot et al. 1998, Pacoureau et al. 2019c), a large abundance of petrels provides easier conditions for skuas to feed themselves and their chicks resulting in a higher breeding success.

Interestingly, we did not find the opposite relationship in the prey dynamics. Our model provided no evidence for a negative effect of the number of skuas on the demographic parameters of the petrel. As skuas prey on both adults and juveniles during the breeding season, we expected a negative effect of the number of skuas on the petrel breeding parameters. This lack of effect could be explained by the large abundance of petrels compared to the skuas on Mayes Island. Oro et al. (2006) reported that in another seabird predator-prey system, the highest breeding success of the prey occurred when the prey/predator ratio was very high. On Mayes Island, the breeding population of petrels is estimated at approximately 142,000 breeding pairs (Barbraud and Delord 2006), and this does not include chicks (around 71,000 each year) and nonbreeders (approximately 30% of the petrels). Hence, there are about 476,000 petrels during a breeding season versus about 200 skuas, resulting in a very high prey/predator ratio. Moreover, Mougeot et al. (1998) showed that skuas breeding at Mayes Island preyed on about 40,000 petrels each breeding season. This corresponds to about 8% of the petrel population of the island. It is therefore possible that skua predation is only a minor factor in shaping petrel demographics, and this effect may be too weak to be detected by our model. Inversely, although the relationships estimated were less clear, our results suggest that the density of skuas tended to increase slightly with the hatching success and breeding success of the prey. However, it is unlikely that the presence of predators increased the reproductive success of petrels. To explain these relationships, we might rely on the other strong effects estimated by our model. Indeed, we found that the number of petrels positively affected the breeding success of skuas and that skuas were sensitive to intraspecific density-dependence. Therefore, years when preys experience a high breeding success correspond to years with particularly abundant food resources for skuas and this until the end of the breeding season. Since skuas are potentially less affected by intraspecific density-dependence than by abundance of prey, they could consequently breed in higher numbers in the breeding area.

### Effects of intraspecific density-dependence

For skuas, we found negative density-dependent effects on breeding success and probability to fledge two chicks, in accordance with our predictions. Egg and chick predation by conspecifics has been reported in the Great Skua (*Catharacta skua*) (Hamer et al. 1991, Ratcliffe and Furness 1999). Hence, a higher abundance of skuas increases the risk of predation on eggs and chicks, resulting in higher breeding failure. To avoid predation by conspecifics, the skuas start defending their territories from conspecifics just a few days after arrival on a breeding site until the end of the season. This activity is energetically costly and may also limit the time spent searching for food, potentially limiting energy investment in reproduction. The heterogeneous habitat hypothesis already demonstrated in territorial birds (Dhondt et al. 1992, Krüger and Lindström 2001, Ferrer and Donazar 2015) could also explain the relationships we found. Indeed, when the population increases, some individuals may be forced to occupy poorer quality habitats, resulting in lower reproductive success. We did not find an effect of density-dependence on the breeding probability of skuas. As skuas are territorial with high site fidelity, we hypothesized that in years with a high abundance of skuas, the breeding probability would decrease, as all the skuas would not succeed in acquiring a territory. It is possible that we did not observe this effect because the logistic function used for density-dependence does not accurately model the territory acquisition dynamics by floaters (e.g. van de Pol et al. 2010, Barraquand et al. 2014).

We estimated that the breeding success of skuas was affected by both predator-prey relationships and intraspecific density-dependence. Predator-prey relationships had a higher contribution to the variability in breeding success of skuas than the density-dependent effect. Hamer et al. (1991) reported that, following a reduction of sandeel (*Ammodytes marinus*) abundance, great skua increased their foraging effort reducing the adult territorial attendance. In turn, breeding failure increased due to predation from adults of neighboring territories. We then may assume that petrel abundance allowed a suitable territorial attendance for skuas reducing the negative density-dependent effects such as chick predation by conspecifics.

For petrels, we found a negative relationship between the breeding success of petrel that did not breed the previous year and the number of petrels on the colony. Combined effects of density-dependence and climate have already been observed in petrels, with a lower winter survival when density is high (Barbraud and Weimerskirch 2003), suggesting a mechanism of competition between conspecifics for food resources. As nonbreeders are known to be in poorer condition than breeders (Chastel et al. 1995), they are potentially more sensitive to the competition for food resources explaining why this effect was only found on petrels that were nonbreeders the previous years. Interestingly, we found a positive intraspecific density-dependence relationship on the breeding probability of petrels that bred the previous year. This suggests that years with a high abundance of petrels reflected a good return rate to the breeding site because environmental conditions were favorable for breeding. This is in agreement with studies showing that petrels might skip breeding and take sabbatical years when environmental conditions are poor (Warham 1990, Chastel et al. 1995).

### Effects of environmental conditions

Breeding probability of petrels tended to be impacted by two of the environmental covariates tested, namely SSTa and Chla. This effect of environmental conditions on the breeding probability is in accordance with previous research showing that the body condition of petrels might impact their decision to attempt breeding (Warham 1990, Chastel et al. 1995). High Chla increases resources availability for organisms at higher trophic levels (macrozooplankton, fishes), which are consumed by petrels (Cherel et al. 2002). Consequently, high Chla may increase abundance of petrel prey, with a positive effect on the breeding performances and body condition of petrels. Unexpectedly, we detected a positive effect of SSTa on breeding probability of petrels. This result is surprising as previous study showed that warm SST events negatively affected the breeding performances and body condition of petrels at Kerguelen Islands (Guinet et al. 1998). Indeed, high SST generally reduces vertical mixing and provides poor growing conditions for zooplankton communities that in turn reduce trophic resources for seabirds (Barbraud et al. 2012, Sydeman et al. 2015). However, it has been showed recently that during the pre-laying period petrels use water masses situated at more northerly latitudes than during the winter period or the breeding period (Quillfeldt et al. 2020), where relationships between SST and primary productivity may differ. Indeed, the covariance between SST and Chla depends on location and shows particularly complex patterns in the Southern Ocean (Dunstan et al. 2018). Positive effects of SSTa have already been identified in other sub-antarctic seabirds (Pinaud and Weimerskirch 2002, Nevoux et al. 2007, Horswill et al. 2014). Furthermore, we estimated that Chla, at the bottom of the trophic food chain, had a higher effect on the breeding probability than SSTa which reflect oceanographic conditions. This indicated that the effect size of environmental covariates increased when the covariates approached the trophic level occupied by the prey of the petrels, suggesting a bottom-up mechanism. This result is consistent with many studies showing that climatic conditions affect seabirds through indirect processes by influencing prey availability and resulting in changes in their dynamics (Frederiksen et al. 2006, Barbraud et al. 2012, Jenouvrier 2013, Lauria et al. 2013).

We did not detect any relationship between the breeding parameters of the skua and the environmental covariates. This lack of effect could be explained by an absence of a direct link between skuas and the environmental covariates tested, as breeding skuas remain on their territory to defend it or to forage. However, we found an effect of SAM on the apparent survival. This effect was detected only for skuas that were nonbreeders during the previous season. It was proposed that only seabirds attaining a threshold condition decide to breed (Weimerskirch 1992), suggesting that nonbreeders are generally in poorer conditions than breeders (Chastel et al. 1995, Cam et al. 1998) and thus more sensitive to environmental conditions. Nevertheless, we found a positive relationship between survival and SAM whereas we expected a negative relationship. Indeed, skuas mainly overwinter north of the polar front (Delord et al. 2018) where positive phases of SAM induce warm SST, low surface Chla concentration (Lovenduski and Gruber 2005), and thus potentially poor feeding conditions. However, only breeding skuas were studied in Delord et al. (2018) and nonbreeding individuals may use different wintering areas where the relationships between SAM and oceanographic variables differ. Several studies reviewed in Jenouvrier (2013) highlighted multifaceted effects of climatic conditions on the demography of seabirds involving direct, time-lagged and non-linear effect, which we did not considered here. Therefore, despite the important contribution of our approach in understanding the effect of the environment in our predator-prey system, disentangling in details the complex mechanisms between environmental covariates and their effects on the demography of the two seabirds remain challenging.

### A bottom-up dynamic in a predator-prey system

Overall, our study has highlighted the important role of bottom-up processes in the dynamics of this marine predator-prey system, *i.e.* the dynamics of these two seabirds was mostly driven by food availability. Petrel dynamics were more strongly affected by environmental covariates near to their trophic level and the number of petrels impacted the dynamics of skuas. The bottom-up control of demographic rates in oceanic predators have been largely assumed (Jenouvrier 2013). This is because the functioning of oceanic systems is controlled and structured by physical processes impacting nutrient fluxes (Behrenfeld et al. 2006) and then the whole trophic food web. We found no evidence of top-down processes, *i.e.* predation effects, in this system, although these two mechanisms have been found to jointly affect ecosystems (Hunter and Price 1992, Sinclair et al. 2003) including other seabird systems (Horswill et al. 2014, 2016, Perkins et al. 2018). Effects of skua predation on petrels were expected, based on their diet during the breeding season. However, given the very large number of petrels present on the island compared to the number of predators, the impact of predation may have been too small to be detected by our model.

## Conclusion

This multispecies IPM framework allowed us to estimate demographic parameters and abundances for both skuas and petrels. Taking into account both species interactions and environmental covariates in the same analysis improved our understanding of species dynamics. We concluded that bottom-up mechanisms are the main drivers of this skua-petrel system. Generalizing such assessments of interspecific relationships and environmental conditions in a single demographic framework may be essential to predict how contrasted climatic scenarios will affect communities. A promising avenue of research in multispecies IPMs lies in fitting models to data on a large number of species, which will much likely require further methodological developments.

## Acknowledgments

This study was made possible thanks to all the field workers involved in the monitoring programs on Brown Skuas and Blue Petrels since 1985 at Mayes Island, Kerguelen Islands. These monitoring programs were supported financially and logistically by the French Polar Institute IPEV (program 109, resp. Henri Weimerskirch), the Zone Atelier Antarctique (CNRS-INEE), Terres Australes et Antarctiques Françaises. All work was carried out in accordance with the guidelines of the IPEV ethics committee. We thank Chloé R. Nater for constructive feedback and helpful suggestions on the manuscript. We acknowledge Dominique Joubert for the management of the demographic database. We thank Dave Koons and an anonymous reviewer for useful comments that helped improved a previous version of the manuscript. This research was funded by the French National Research Agency (grant ANR-16-CE02-0007).

