## Appendix S1 for "Multispecies integrated population model reveals bottom-up dynamics in a seabird predator-prey system"

### Appendix S1: Description of state variables, demographic parameters and notations

Table S1: Description of all state variables, parameters and notations.

| Parameters | Definition |
| --- | --- |
| **State variable** | |
| $N_{J1}$ | Estimated number of individuals in their first year i.e. between 0 and 1 year old. Indexed by species and time. Ex: $N_{J1,S,t}$ |
| $N_{J2}$ | Estimated number of second year individuals i.e. between 1 and 2 years old. Indexed by species and time. Ex: $N_{J2,S,t}$ |
| $N_{J3}$ | Estimated number of third year individuals i.e. between 2 and 3 years old. Indexed by species and time. Ex: $N_{J3,S,t}$ |
| $N_{J4}$ | Estimated number of fourth year individuals i.e. between 3 and 4 years old. Indexed by species and time. Ex: $N_{J4,S,t}$ |
| $N_{J4NB}$ | Estimated number of individuals of 4 years old and older that did not attempt to breed. Indexed by species and time. Ex: $N_{J4NB,S,t}$ |
| $N_{J4B}$ | Estimated number of individuals of 4 years old and older that attempted to breed for the first time. Indexed by species and time. Ex: $N_{J4B,S,t}$ |
| $N_{J4+}$ | Estimated number of individuals of 4 years and older. Indexed by species and time. Ex: $N_{J4+,S,t}$ |
| $N_{alive}$ | Estimated number of surviving adults. Indexed by species and time. Ex: $N_{alive,S,t}$ |
| $N_{Balive}$ | Estimated number of surviving adults that attempted to breed. Indexed by species and time. Ex: $N_{Balive,S,t}$ |
| $N_{B}$ | Estimated total number of individuals that attempted to breed. Indexed by species and time. Ex: $N_{B,S,t}$ |
| $N_{NB}$ | Estimated number of surviving adults that did not attempted to breed. Indexed by species and time. Ex: $N_{NB,S,t}$ |
| $N_{adtot}$ | Estimated total number of individuals. Indexed by species and time. Ex: $N_{adtot,S,t}$ |
| $N_{FB}$ | Estimated number of failed breeders. Only for skuas. Indexed by species and time. Ex: $N_{FB,S,t}$ |
| $N_{FBE}$ | Estimated number of failed breeders at the stage egg. Only for petrels. Indexed by species and time. Ex: $N_{FBE,P,t}$ |
| $N_{SH}$ | Estimated number of individuals with an egg successfully hatched. Only for petrels. Indexed by species and time. Ex: $N_{SH,P,t}$ |
| $N_{FBC}$ | Estimated number of failed breeders at the chick stage. Only for petrels. Indexed by species and time. Ex: $N_{FBC,P,t}$ |
| $N_{SB}$ | Estimated number of successful breeders. Indexed by species and time. Ex : $N_{SB,S,t}$ |
| $N_{SB1}$ | Estimated number of successful breeders with one chick. Only for skuas. Indexed by species and time. Ex: $N_{SB1,S,t}$ |
| $N_{SB2}$ | Estimated number of successful breeders with two chicks. Only for skuas. Indexed by species and time. Ex: $N_{SB2,S,t}$ |
| **Demographic parameters** | |
| $\phi_{J1}$ | Survival of $N_{J1}$ individuals. Indexed by species. Ex: $\phi_{J1,S}$ |
| $\phi_{J2}$ | Survival of $N_{J2}$ individuals. Indexed by species. Ex: $\phi_{J2,S}$ |
| $\phi_{J3}$ | Survival of $N_{J3}$ individuals. Indexed by species. Ex: $\phi_{J3,S}$ |
| $\phi_{J4}$ | Survival of $N_{J4+}$ individuals. Indexed by species. Ex: $\phi_{J4,S}$ |
| $Pr$ | First breeding attempt probability. Indexed by species and time. Ex:${Pr}_{S,t}$ |
| $\phi$ | Adult apparent survival. Indexed by species, previous breeding status and time. Ex: $\phi_{S,\tilde{B},t}$ |
| $\beta$ | Breeding probability. Indexed by species, previous breeding status and time. Ex: $\beta_{S,\tilde{B},t}$ |
| $\omega$ | Hatching success probability. Only for petrels. Indexed by species, previous breeding status and time. Ex: $\omega_{P,\tilde{B},t}$ |
| $\gamma$ | Breeding success probability. Indexed by species, previous breeding status and time. Ex: $\gamma_{S,\tilde{B},t}$ |
| $\delta$ | Breeding success with two chicks probability. Only for skuas. Indexed by species, previous breeding status and time. Ex: $\delta_{S,\tilde{B},t}$ |
| **Data** | |
| $Y$ | Number of territories/burrows occupied multiplied by 2, corresponding to the total number of adults in the monitoring area. Indexed by species and time. Ex: $Y_{S,t}$ |
| $Ch$ | Capture history. Indexed by species. Ex: ${Ch}_{S}$ |
| $f$ | Fecundity. Fixed values: 1 or 2 chicks per pair of seabirds. Indexed by breeding status and species. Ex: $f_{SB1,S}$ |
| $N_{im}$ | Number of immigrant individuals, *i.e.* individuals newly marked that came to the monitoring area to breed. Fixed values. Indexed by species and time. Ex: $N_{im,S,t}$. |
| $SAM$ | Environmental covariate: Southern Annular Mode index. Indexed by demographic parameter, species and time. Ex: ${SAM}_{\gamma,S,t}$ |
| $SSTa$ | Environmental covariate: Sea Surface Temperature anomalies. Only for petrels. Indexed by demographic parameter, species and time. Ex: ${SST}_{\gamma,P,t}$ |
| $Chla$ | Environmental covariate: Chlorophyll a concentration. Only for petrels. Indexed by demographic parameter, species and time. Ex: ${Chla}_{\gamma,P,t}$ |
| **Logit-linear relationships** | |
| $\mu$ | Intercept of the logit-linear regressions on demographic parameters. Indexed by demographic parameter, species and previous breeding status. Ex: $\mu_{\gamma,S,\tilde{B}}$ |
| $\alpha$ | Slope on the effect tested (environmental covariate, density-dependence or predator-prey relationship) in the logit linear regressions on the demographic parameters. Indexed by the effect tested, demographic parameter, species and previous breeding status. Ex: $\alpha_{PP,\gamma,S,\tilde{B}}$ |
| $\varepsilon$ | Yearly random effect in the logit linear regressions on the demographic parameters. Indexed by the demographic parameter, species, previous breeding status and time. Ex: $\varepsilon_{\gamma,S,\tilde{B},t}$ |
| ${\sigma^{2}}_{\varepsilon}$ | Variance of the yearly random effect. Indexed by the demographic parameter, species and previous breeding status. Ex: ${\sigma^{2}}_{\varepsilon,\gamma,S,\tilde{B}}$ |
| **Breeding status** | |
| *NB* | Non breeder. |
| *FB* | Failed breeder. Only for skuas. |
| *FBE* | Failed breeder at the stage egg. Only for petrels. |
| *FBC* | Failed breeder at the stage chick. Only for petrels. |
| *SB* | Successful breeder. Only for petrels. |
| *SB1* | Successful breeder with one chick. Only for skuas. |
| *SB2* | Successful breeder with two chicks. Only for skuas. |
| *C* | Uncertain. |
| **Index** | |
| *t* | Time |
| *S* | Brown Skua |
| *P* | Blue Petrel |
| *X* | S or P |
| $\tilde{B}$ | Breeder during the previous breeding season |
| $\tilde{NB}$ | Non breeder during the previous breeding season |
| *PP* | Predator-Prey |
| $DD$ | Intraspecific Density-Dependence |
