## Appendix S2 for "Multispecies integrated population model reveals bottom-up dynamics in a seabird predator-prey system"

### Appendix S2: Logit-linear regression equations to estimate the effects of covariates and state variables on demographic parameters

#### Brown Skua demographic parameters

##### Survival

We modelled the adult apparent survival probability for skuas that bred the previous year ${(\phi}_{S,\tilde{B}})$ with a logit link:

$logit{(\phi}_{S,\tilde{B},t-1})= \mu_{\phi,S,\tilde{B}}+ \alpha_{SAM,\phi,S,\tilde{B}} \times{SAM}_{\phi,S,t}+\alpha_{DD,\phi,S,\tilde{B}}\times N_{adtot,S,t-1}+ \varepsilon_{\phi,S,\tilde{B},t}$

$\varepsilon_{\phi,S,\tilde{B},t} \sim N(0, {\sigma^{2}}_{\varepsilon,\phi,S,\tilde{B}})$	(1)

with $\mu_{\phi,S,\tilde{B}}$ the intercept, $\alpha_{SAM,\phi,S,\tilde{B}}$the slope for the climatic covariate ${SAM}_{\phi,S},$ $\alpha_{DD,\phi,S,\tilde{B}}$the slope indicating the strength of the intraspecific density-dependence with $N_{adtot,S}$the number of adult skuas, $\varepsilon_{\phi,S,\tilde{B}}$a yearly random effect and ${\sigma^{2}}_{\varepsilon,\phi,S,\tilde{B}}$ its temporal variance.

We modelled the adult apparent survival probability for skuas that did not breed the previous year${(\phi}_{S,\tilde{NB}}$) with a logit link:

$logit{(\phi}_{S,\tilde{NB},t-1})= \mu_{\phi,S,\tilde{NB}}+ \alpha_{SAM,\phi,S,\tilde{NB}} \times{SAM}_{\phi,S,t}+\alpha_{DD,\phi,S,\tilde{NB}}\times N_{adtot,S,t-1}+ \varepsilon_{\phi,S,\tilde{NB},t}$

$\varepsilon_{\phi,S,\tilde{NB},t} \sim N(0, {\sigma^{2}}_{\varepsilon,\phi,S,\tilde{NB}})$	(2)

with $\mu_{\phi,S,\tilde{NB}}$the intercept, $\alpha_{SAM,\phi,S,\tilde{NB}}$the slope for the climatic covariate ${SAM}_{\phi,S}$, $\alpha_{DD,\phi,S,\tilde{NB}}$ the slope indicating the strength of the intraspecific density-dependence with $N_{adtot,S}$the number of adult skuas, $\varepsilon_{\phi,S,\tilde{NB}}$a yearly random effect and ${\sigma^{2}}_{\varepsilon,\phi,S,\tilde{NB}}$its temporal variance.

##### Breeding probability

We modelled the breeding probability for skuas that bred the previous year ${(\beta}_{S,\tilde{B}})$ with a logit link:

$logit{(\beta}_{S,\tilde{B},t-1})= \mu_{\beta,S,\tilde{B}}+ \alpha_{SAM,\beta,S,\tilde{B}} \times{SAM}_{\beta,S,t}+\alpha_{DD,\beta,S,\tilde{B}}\times N_{adtot,S,t}+ \varepsilon_{\beta,S,\tilde{B},t}$

$\varepsilon_{\beta,S,\tilde{B},t} \sim N(0, {\sigma^{2}}_{\varepsilon,\beta,S,\tilde{B}})$	(3)

with $\mu_{\beta,S,\tilde{B}}$the intercept, $\alpha_{SAM,\beta,S,\tilde{B}}$the slope for the climatic covariate ${SAM}_{\beta,S}$, $\alpha_{DD,\beta,S,\tilde{B}}$ the slope indicating the strength of the intraspecific density-dependence with $N_{adtot,S}$the number of adult skuas, $\varepsilon_{\beta,S,\tilde{B}}$ a yearly random effect and ${\sigma^{2}}_{\varepsilon,\beta,S,\tilde{B}}$its temporal variance.

We modelled the breeding probability for skuas that did not breed the previous year ${(\beta}_{S,\tilde{NB}})$ with a logit link:

$logit{(\beta}_{S,\tilde{NB},t-1})= \mu_{\beta,S,\tilde{NB}}+ \alpha_{SAM,\beta,S,\tilde{NB}} \times{SAM}_{\beta,S,t}+\alpha_{DD,\beta,S,\tilde{NB}}\times N_{adtot,S,t}+ \varepsilon_{\beta,S,\tilde{NB},t}$

$\varepsilon_{\beta,S,\tilde{NB},t} \sim N(0,{\sigma^{2}}_{\varepsilon,\beta,S,\tilde{NB}})$	(4)

with $\mu_{\beta,S,\tilde{NB}}$ the intercept, $\alpha_{SAM,\beta,S,\tilde{NB}}$the slope for the climatic covariate ${SAM}_{\beta,S}$, $\alpha_{DD,\beta,S,\tilde{NB}}$ the slope indicating the strength of the intraspecific density-dependence with $N_{adtot,S}$the number of adult skuas, $\varepsilon_{\beta,S,\tilde{NB}}$ a yearly random effect and ${\sigma^{2}}_{\varepsilon,\beta,S,\tilde{NB}}$ its temporal variance.

##### Breeding success

We modelled the breeding success probability for skuas that bred the previous year ${(\gamma}_{S,\tilde{B}})$ with a logit link:

$logit{(\gamma}_{S,\tilde{B},t-1})= \mu_{\gamma,S,\tilde{B}}+ \alpha_{SAM,\gamma,S,\tilde{B}} \times{SAM}_{\gamma,S,t}+\alpha_{DD,\gamma,S,\tilde{B}}\times N_{adtot,S,t}+\alpha_{PP,\gamma,S,\tilde{B}}\times N_{adtot,P,t}+ \varepsilon_{\gamma,S,\tilde{B},t}$

$\varepsilon_{\gamma,S,\tilde{B},t} \sim N(0, {\sigma^{2}}_{\varepsilon,\gamma,S,\tilde{B}})$	(5)

with $\mu_{\gamma,S,\tilde{B}}$ the intercept, $\alpha_{SAM,\gamma,S,\tilde{B}}$the slope for the climatic covariate ${SAM}_{\gamma,S}$ ,$\alpha_{DD,\gamma,S,\tilde{B}}$the slope indicating the strength of the intraspecific density-dependence with $N_{adtot,S}$the number of adult skuas, $\alpha_{PP,\gamma,S,\tilde{B}}$ the slope indicating the strength of the predator-prey relationship with $N_{adtot,P}$the number of adult petrels, $\varepsilon_{\gamma,S,\tilde{B},t}$ a yearly random effect and ${\sigma^{2}}_{\varepsilon,\gamma,S,\tilde{B}}$ its temporal variance.

We modelled the breeding success probability for skuas that did not breed the previous year ${(\gamma}_{S,\tilde{NB}})$ with a logit link:

$logit{(\gamma}_{S,\tilde{NB},t-1})= \mu_{\gamma,S,\tilde{NB}}+ \alpha_{SAM,\gamma,S,\tilde{NB}} \times{SAM}_{\gamma,S,t}+\alpha_{DD,\gamma,S,\tilde{NB}}\times N_{adtot,S,t}+\alpha_{PP,\gamma,S,\tilde{NB}}\times N_{adtot,P,t}+ \varepsilon_{\gamma,S,\tilde{NB},t}$

$\varepsilon_{\gamma,S,\tilde{NB},t} \sim N(0, {\sigma^{2}}_{\varepsilon,\gamma,S,\tilde{NB}})$	(6)

with $\mu_{\gamma,S,\tilde{NB}}$ the intercept, $\alpha_{SAM,\gamma,S,\tilde{NB}}$the slope for the climatic covariate ${SAM}_{\gamma,S}$ ,$\alpha_{DD,\gamma,S,\tilde{NB}}$the slope indicating the strength of the intraspecific density-dependence with $N_{adtot,S}$the number of adult skuas, $\alpha_{PP,\gamma,S,\tilde{NB}}$ the slope indicating the strength of the predator-prey relationship with $N_{adtot,P}$the number of adult petrels, $\varepsilon_{\gamma,S,\tilde{NB},t}$ a yearly random effect and ${\sigma^{2}}_{\varepsilon,\gamma,S,\tilde{NB}}$ its temporal variance.

##### Breeding success with two chicks

We modelled the breeding success probability with two chicks for skuas that bred the previous year ${(\delta}_{S,\tilde{B}})$ with a logit link:

$logit{(\delta}_{S,\tilde{B},t-1})= \mu_{\delta,S,\tilde{B}}+ \alpha_{SAM,\delta,S,\tilde{B}} \times{SAM}_{\delta,S,t}+\alpha_{DD,\delta,S,\tilde{B}}\times N_{adtot,S,t}+\alpha_{PP,\delta,S,\tilde{B}}\times N_{adtot,P,t}+ \varepsilon_{\delta,S,\tilde{B},t}$

$\varepsilon_{\delta,S,\tilde{B},t} \sim N(0, {\sigma^{2}}_{\varepsilon,\delta,S,\tilde{B}})$	(7)

with $\mu_{\delta,S,\tilde{B}}$ the intercept, $\alpha_{SAM,\delta,S,\tilde{B}}$ the slope for the climatic covariate ${SAM}_{\delta,S}$, $\alpha_{DD,\delta,S,\tilde{B}}$the slope indicating the strength of the intraspecific density-dependence with $N_{adtot,S}$the number of adult skuas, $\alpha_{PP,\delta,S,\tilde{B}}$the slope indicating the strength of the predator-prey relationship with $N_{adtot,P}$the number of adult petrels, $\varepsilon_{\delta,S,\tilde{B}}$ a yearly random effect and ${\sigma^{2}}_{\varepsilon,\delta,S,\tilde{B}}$ its temporal variance.

We modelled the breeding success probability with two chicks for skuas that did not breed the previous year ${(\delta}_{S,\tilde{NB}})$ with a logit link:

$logit{(\delta}_{S,\tilde{NB},t-1})= \mu_{\delta,S,\tilde{NB}}+ \alpha_{SAM,\delta,S,\tilde{NB}} \times{SAM}_{\delta,S,t}+\alpha_{DD,\delta,S,\tilde{NB}}\times N_{adtot,S,t}+\alpha_{PP,\delta,S,\tilde{NB}}\times N_{adtot,P,t}+ \varepsilon_{\delta,S,\tilde{NB},t}$

$\varepsilon_{\delta,S,\tilde{NB},t} \sim N(0, {\sigma^{2}}_{\varepsilon,\delta,S,\tilde{NB}})$	(8)

with $\mu_{\delta,S,\tilde{NB}}$ the intercept, $\alpha_{SAM,\delta,S,\tilde{NB}}$the slope for the climatic covariate ${SAM}_{\delta,S}$, $\alpha_{DD,\delta,S,\tilde{NB}}$the slope indicating the strength of the intraspecific density-dependence with $N_{adtot,S}$the number of adult skuas, $\alpha_{PP,\delta,S,\tilde{NB}}$the slope indicating the strength of the predator-prey relationship with $N_{adtot,P}$the number of adult petrels, $\varepsilon_{\delta,S,\tilde{NB}}$ a yearly random effect and ${\sigma^{2}}_{\varepsilon,\delta,S,\tilde{NB}}$ its temporal variance.

#### Blue Petrel demographic parameters

##### Survival

We modelled the adult apparent survival probability for petrels that bred the previous year ${(\phi}_{P,\tilde{B}})$ with a logit link:

$logit{(\phi}_{P,\tilde{B},t-1})= \mu_{\phi,P,\tilde{B}}+ \alpha_{SAM,\phi,P,\tilde{B}} \times{SAM}_{\phi,P,t}+ \alpha_{SSTa,\phi,P,\tilde{B}} \times{SSTa}_{\phi,P,t}+ \alpha_{Chla,\phi,P,\tilde{B}} \times{Chla}_{\phi,P,t}+\alpha_{DD,\phi,P,\tilde{B}}\times N_{adtot,P,t-1}+\alpha_{PP,\phi,P,\tilde{B}}\times N_{adtot,S,t-1}+ \varepsilon_{\phi,P,\tilde{B},t}$

$\varepsilon_{\phi,P,\tilde{B},t} \sim N(0, {\sigma^{2}}_{\varepsilon,\phi,P,\tilde{B}})$	(9)

with $\mu_{\phi,P,\tilde{B}}$ the intercept, $\alpha_{SAM,\phi,P,\tilde{B}}$the slope for the climatic covariate ${SAM}_{\phi,P}$, $\alpha_{SSTa,\phi,P,\tilde{B}}$ the slope for the climatic covariate${SSTa}_{\phi,P}$, $\alpha_{Chla,\phi,P,\tilde{B}}$the slope for the climatic covariate${Chla}_{\phi,P}$, $\alpha_{DD,\phi,P,\tilde{B}}$the slope indicating the strength of the intraspecific density-dependence with $N_{adtot,P}$the number of adult petrels, $\alpha_{PP,\phi,P,\tilde{B}}$ the slope indicating the strength of the predator-prey relationship with $N_{adtot,S}$the number of adult skuas, $\varepsilon_{\phi,P,\tilde{B}}$ a yearly random effect and ${\sigma^{2}}_{\varepsilon,\phi,P,\tilde{B}}$ its temporal variance.

We modelled the adult apparent survival probability for petrels that did not breed the previous year ${(\phi}_{P,\tilde{NB}})$ with a logit link:

$logit{(\phi}_{P,\tilde{NB},t-1})= \mu_{\phi,P,\tilde{NB}}+ \alpha_{SAM,\phi,P,\tilde{NB}} \times{SAM}_{\phi,P,t}+ \alpha_{SSTa,\phi,P,\tilde{NB}} \times{SSTa}_{\phi,P,t}+ \alpha_{Chla,\phi,P,\tilde{NB}} \times{Chla}_{\phi,P,t}+\alpha_{DD,\phi,P,\tilde{NB}}\times N_{adtot,P,t}+\alpha_{PP,\phi,P,\tilde{NB}}\times N_{adtot,S,t}+ \varepsilon_{\phi,P,\tilde{NB},t}$

$\varepsilon_{\phi,P,\tilde{NB},t} \sim N(0, {\sigma^{2}}_{\varepsilon,\phi,P,\tilde{NB}})$	(10)

with $\mu_{\phi,P,\tilde{NB}}$ the intercept, $\alpha_{SAM,\phi,P,\tilde{NB}}$the slope for the climatic covariate ${SAM}_{\phi,P}$, $\alpha_{SSTa,\phi,P,\tilde{NB}}$the slope for the climatic covariate${SSTa}_{\phi,P}$, $\alpha_{Chla,\phi,P,\tilde{NB}}$the slope for the climatic covariate${Chla}_{\phi,P}$, $\alpha_{DD,\phi,P,\tilde{NB}}$the slope indicating the strength of the intraspecific density-dependence with $N_{adtot,P}$the number of adult petrels, $\alpha_{PP,\phi,P,\tilde{NB}}$the slope indicating the strength of the predator-prey relationship with $N_{adtot,S}$the number of adult skuas, $\varepsilon_{\phi,P,\tilde{NB}}$ a yearly random effect and ${\sigma^{2}}_{\varepsilon,\phi,P,\tilde{NB}}$ its temporal variance.

##### Breeding probability

We modelled the breeding probability for petrels that bred the previous year ${(\beta}_{P,\tilde{B}})$ with a logit link:

$logit{(\beta}_{P,\tilde{B},t-1})= \mu_{\beta,P,\tilde{B}}+ \alpha_{SAM,\beta,P,\tilde{B}} \times{SAM}_{\beta,P,t}+ \alpha_{SSTa,\beta,P,\tilde{B}} \times{SSTa}_{\beta,P,t}+ \alpha_{Chla,\beta,P,\tilde{B}} \times{Chla}_{\beta,P,t}+\alpha_{DD,\beta,P,\tilde{B}}\times N_{adtot,P,t}+ \varepsilon_{\beta,P,\tilde{B},t}$

$\varepsilon_{\beta,P,\tilde{B},t} \sim N(0, {\sigma^{2}}_{\varepsilon,\beta,P,\tilde{B}})$	(11)

with $\mu_{\beta,P,\tilde{B}}$ the intercept, $\alpha_{SAM,\beta,P,\tilde{B}}$the slope for the climatic covariate${SAM}_{\beta,P,t}$, $\alpha_{SSTa,\beta,P,\tilde{B}}$the slope for the climatic covariate${SSTa}_{\beta,P}$, $\alpha_{Chla,\beta,P,\tilde{B}}$the slope for the climatic covariate${Chla}_{\beta,P}$, $\alpha_{DD,\beta,P,\tilde{B}}$the slope indicating the strength of the intraspecific density-dependence with $N_{adtot,P}$the number of adult petrels, $\varepsilon_{\beta,P,\tilde{B}}$ a yearly random effect and ${\sigma^{2}}_{\varepsilon,\beta,P,\tilde{B}}$ its temporal variance.

We modelled the breeding probability for petrels that did not breed the previous year ${(\beta}_{P,\tilde{NB}})$ with a logit link:

$logit{(\beta}_{P,\tilde{NB},t-1})= \mu_{\beta,P,\tilde{NB}}+ \alpha_{SAM,\beta,P,\tilde{NB}} \times{SAM}_{\beta,P,t}+ \alpha_{SSTa,\beta,P,\tilde{NB}} \times{SSTa}_{\beta,P,t}+ \alpha_{Chla,\beta,P,\tilde{NB}} \times{Chla}_{\beta,P,t}+\alpha_{DD,\beta,P,\tilde{NB}}\times N_{adtot,P,t}+ \varepsilon_{\beta,P,\tilde{NB},t}$

$\varepsilon_{\beta,P,\tilde{NB},t} \sim N(0, {\sigma^{2}}_{\varepsilon,\beta,P,\tilde{NB}})$	(12)

with $\mu_{\beta,P,\tilde{NB}}$ the intercept, $\alpha_{SAM,\beta,P,\tilde{NB}}$the slope for the climatic covariate${SAM}_{\beta,P}$, $\alpha_{SSTa,\beta,P,\tilde{NB}}$the slope for the climatic covariate${SSTa}_{\beta,P}$, $\alpha_{Chla,\beta,P,\tilde{NB}}$the slope for the climatic covariate${Chla}_{\beta,P}$, $\alpha_{DD,\beta,P,\tilde{NB}}$the slope indicating the strength of the intraspecific density-dependence with $N_{adtot,P}$the number of adult petrels, $\varepsilon_{\beta,P,\tilde{NB}}$a yearly random effect and ${\sigma^{2}}_{\varepsilon,\beta,P,\tilde{NB}}$ its temporal variance.

##### Hatching probability

We modelled the hatching probability for petrels that bred the previous year ${(\omega}_{P,\tilde{B}})$ with a logit link:

$logit{(\omega}_{P,\tilde{B},t-1})= \mu_{\omega,P,\tilde{B}}+ \alpha_{SAM,\omega,P,\tilde{B}} \times{SAM}_{\omega,P,t}+ \alpha_{SSTa,\omega,P,\tilde{B}} \times{SSTa}_{\omega,P,t}+ \alpha_{Chla,\omega,P,\tilde{B}} \times{Chla}_{\omega,P,t}+\alpha_{DD,\omega,P,\tilde{B}}\times N_{adtot,P,t}+\alpha_{PP,\omega,P,\tilde{B}}\times N_{adtot,S,t}+ \varepsilon_{\omega,P,\tilde{B},t}$

$\varepsilon_{\omega,P,\tilde{B},t} \sim N(0, {\sigma^{2}}_{\varepsilon,\omega,P,\tilde{B}})$	(13)

with $\mu_{\omega,P,\tilde{B}}$ the intercept, $\alpha_{SAM,\omega,P,\tilde{B}}$the slope for the climatic covariate${SAM}_{\omega,P}$, $\alpha_{SSTa,\omega,P,\tilde{B}}$ the slope for the climatic covariate${SSTa}_{\omega,P}$, $\alpha_{Chla,\omega,P,\tilde{B}}$ the slope for the climatic covariate${Chla}_{\omega,P}$, $\alpha_{DD,\omega,P,\tilde{B}}$ the slope indicating the strength of the intraspecific density-dependence with $N_{adtot,P}$the number of adult petrels, $\alpha_{PP,\omega,P,\tilde{B}}$ the slope indicating the strength of the predator-prey relationship with $N_{adtot,S}$the number of adult skuas, $\varepsilon_{\omega,P,\tilde{B}}$ a yearly random effect and ${\sigma^{2}}_{\varepsilon,\omega,P,\tilde{B}}$ its temporal variance.

We modelled the hatching probability for petrels that did not breed the previous year ${(\omega}_{P,\tilde{NB}})$ with a logit link:

$logit{(\omega}_{P,\tilde{NB}})= \mu_{\omega,P,\tilde{NB}}+ \alpha_{SAM,\omega,P,\tilde{NB}} \times{SAM}_{\omega,P,t}+ \alpha_{SSTa,\omega,P,\tilde{NB}} \times{SSTa}_{\omega,P,t}+ \alpha_{Chla,\omega,P,\tilde{NB}} \times{Chla}_{\omega,P,t}+\alpha_{DD,\omega,P,\tilde{NB}}\times N_{adtot,P,t}+\alpha_{PP,\omega,P,\tilde{NB}}\times N_{adtot,S,t}+ \varepsilon_{\omega,P,\tilde{NB},t}$

$\varepsilon_{\omega,P,\tilde{NB},t} \sim N(0, {\sigma^{2}}_{\varepsilon,\omega,P,\tilde{NB}})$	(14)

with $\mu_{\omega,P,\tilde{NB}}$ the intercept, $\alpha_{SAM,\omega,P,\tilde{NB}}$the slope for the climatic covariate${SAM}_{\omega,P}$, $\alpha_{SSTa,\omega,P,\tilde{NB}}$ the slope for the climatic covariate${SSTa}_{\omega,P}$, $\alpha_{Chla,\omega,P,\tilde{NB}}$ the slope for the climatic covariate${Chla}_{\omega,P}$ , $\alpha_{DD,\omega,P,\tilde{NB}}$ the slope indicating the strength of the intraspecific density-dependence with $N_{adtot,P}$the number of adult petrels, $\alpha_{PP,\omega,P,\tilde{NB}}$ the slope indicating the strength of the predator-prey relationship with $N_{adtot,S}$the number of adult skuas, $\varepsilon_{\omega,P,\tilde{NB}}$ a yearly random effect and ${\sigma^{2}}_{\varepsilon,\omega,P,\tilde{NB}}$ its temporal variance.

##### Breeding success

We modelled the breeding success probability for petrels that bred the previous year ${(\gamma}_{P,\tilde{B}})$ with a logit link:

$logit{(\gamma}_{P,\tilde{B},t-1})= \mu_{\gamma,P,\tilde{B}}+ \alpha_{SAM,\gamma,P,\tilde{B}} \times{SAM}_{\gamma,P,t}+ \alpha_{SSTa,\gamma,P,\tilde{B}} \times{SSTa}_{\gamma,P,t}+ \alpha_{Chla,\gamma,P,\tilde{B}} \times{Chla}_{\gamma,P,t}+\alpha_{DD,\gamma,P,\tilde{B}}\times N_{adtot,P,t}+\alpha_{PP,\gamma,P,\tilde{B}}\times N_{adtot,S,t}+ \varepsilon_{\gamma,P,\tilde{B},t}$

$\varepsilon_{\gamma,P,\tilde{B},t} \sim N(0, {\sigma^{2}}_{\varepsilon,\gamma,P,\tilde{B}})$	(15)

with $\mu_{\gamma,P,\tilde{B}}$ the intercept, $\alpha_{SAM,\gamma,P,\tilde{B}}$the slope for the climatic covariate${SAM}_{\gamma,P}$ , $\alpha_{SSTa,\gamma,P,\tilde{B}}$ the slope for the climatic covariate${SSTa}_{\gamma,P}$, $\alpha_{Chla,\gamma,P,\tilde{B}}$ the slope for the climatic covariate${Chla}_{\gamma,P}$, $\alpha_{DD,\gamma,P,\tilde{B}}$ the slope indicating the strength of the intraspecific density-dependence with $N_{adtot,P}$the number of adult petrels, $\alpha_{PP,\gamma,P,\tilde{B}}$ the slope indicating the strength of the predator-prey relationship with $N_{adtot,S}$the number of adult skuas, $\varepsilon_{\gamma,P,\tilde{B}}$ a yearly random effect and ${\sigma^{2}}_{\varepsilon,\gamma,P,\tilde{B}}$ its temporal variance.

We modelled the breeding success probability for petrels that did not breed the previous year ${(\gamma}_{P,\tilde{NB}})$ with a logit link:

$logit{(\gamma}_{P,\tilde{NB},t-1})= \mu_{\gamma,P,\tilde{NB}}+ \alpha_{SAM,\gamma,P,\tilde{NB}} \times{SAM}_{\gamma,P,t}+ \alpha_{SSTa,\gamma,P,\tilde{NB}} \times{SSTa}_{\gamma,P,t}+ \alpha_{Chla,\gamma,P,\tilde{NB}} \times{Chla}_{\gamma,P,t}+\alpha_{DD,\gamma,P,\tilde{NB}}\times N_{adtot,P,t}+\alpha_{PP,\gamma,P,\tilde{NB}}\times N_{adtot,S,t}+ \varepsilon_{\gamma,P,\tilde{NB},t}$

$\varepsilon_{\gamma,P,\tilde{NB},t} \sim N(0, {\sigma^{2}}_{\varepsilon,\gamma,P,\tilde{NB}})$	(16)

with $\mu_{\gamma,P,\tilde{NB}}$ the intercept, $\alpha_{SAM,\gamma,P,\tilde{NB}}$the slope for the climatic covariate ${SAM}_{\gamma,P}$ , $\alpha_{SSTa,\gamma,P,\tilde{NB}}$the slope for the climatic covariate${SSTa}_{\gamma,P}$, $\alpha_{Chla,\gamma,P,\tilde{NB}}$the slope for the climatic covariate${Chla}_{\gamma,P}$, $\alpha_{DD,\gamma,P,\tilde{NB}}$the slope indicating the strength of the intraspecific density-dependence with $N_{adtot,P}$the number of adult petrels, $\alpha_{PP,\gamma,P,\tilde{NB}}$the slope indicating the strength of the predator-prey relationship with $N_{adtot,S}$the number of adult skuas, $\varepsilon_{\gamma,P,\tilde{NB}}$a yearly random effect and ${\sigma^{2}}_{\varepsilon,\gamma,P,\tilde{NB}}$ its temporal variance.
