## Appendix S3 for "Multispecies integrated population model reveals bottom-up dynamics in a seabird predator-prey system"

### Appendix S3: Demographic parameters and population size estimates

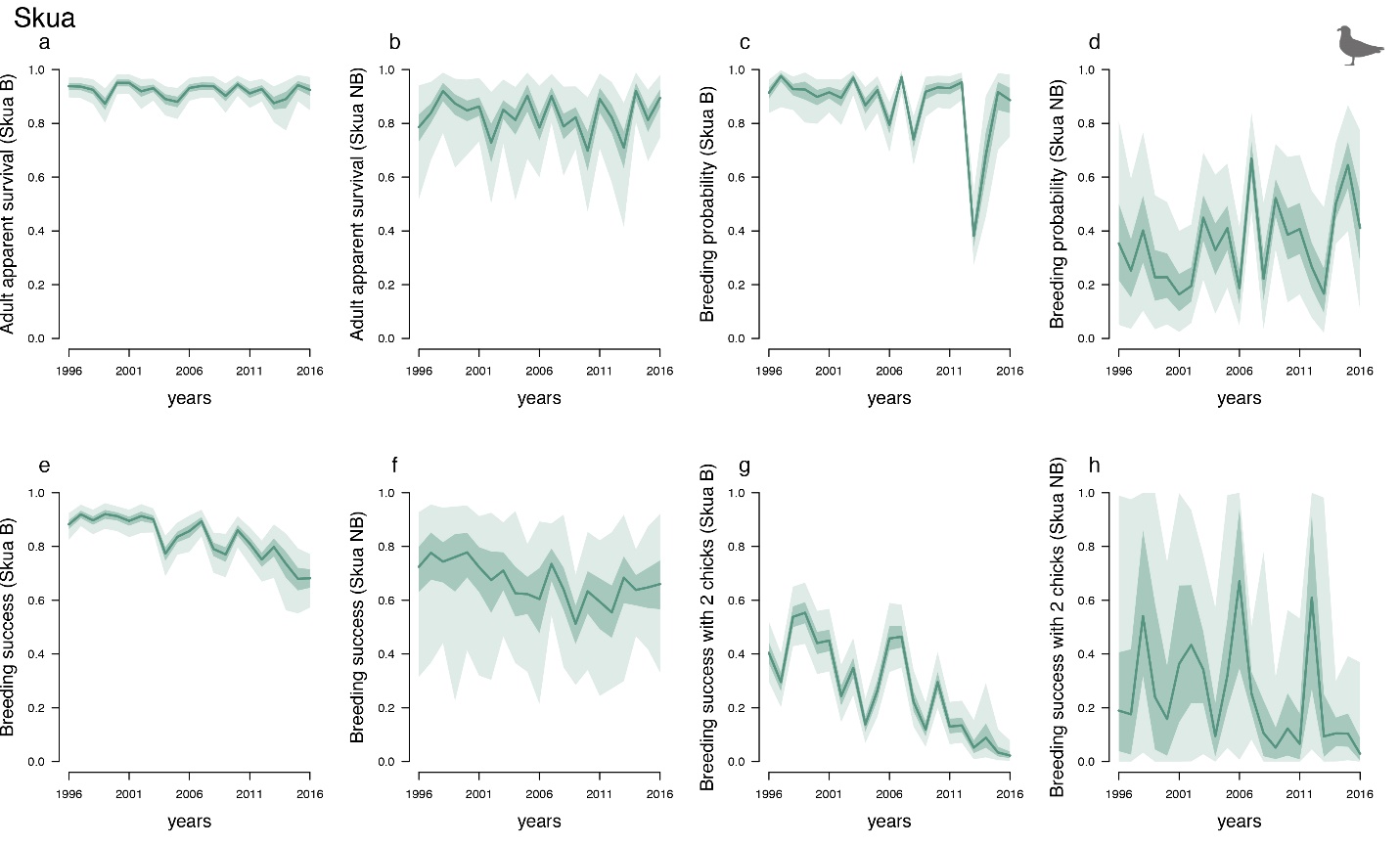

Figure S1: Demographic parameter estimates for each year (1996-2016) for Brown Skuas. Solid lines represent the means of marginal posterior distributions. Shaded areas are the 50% and 95% credibility intervals. (a, b) Estimated adult apparent survival probability$\phi_{S,\tilde{B}}$ and$\phi_{S,\tilde{NB}}$ (c, d) Estimated breeding probability$\beta_{S,\tilde{B}}$ and$\beta_{S,\tilde{NB}}$ (e, f) Estimated breeding success probability$\gamma_{S,\tilde{B}}$ and $\gamma_{S,\tilde{NB}}$(g, h) Estimated breeding success probability for two chicks$\delta_{S,\tilde{B}}$ and $\delta_{S,\tilde{NB}}$ for skuas that have bred ($\tilde{B}$) or have not bred ($\tilde{NB}$), respectively, during the previous breeding season.

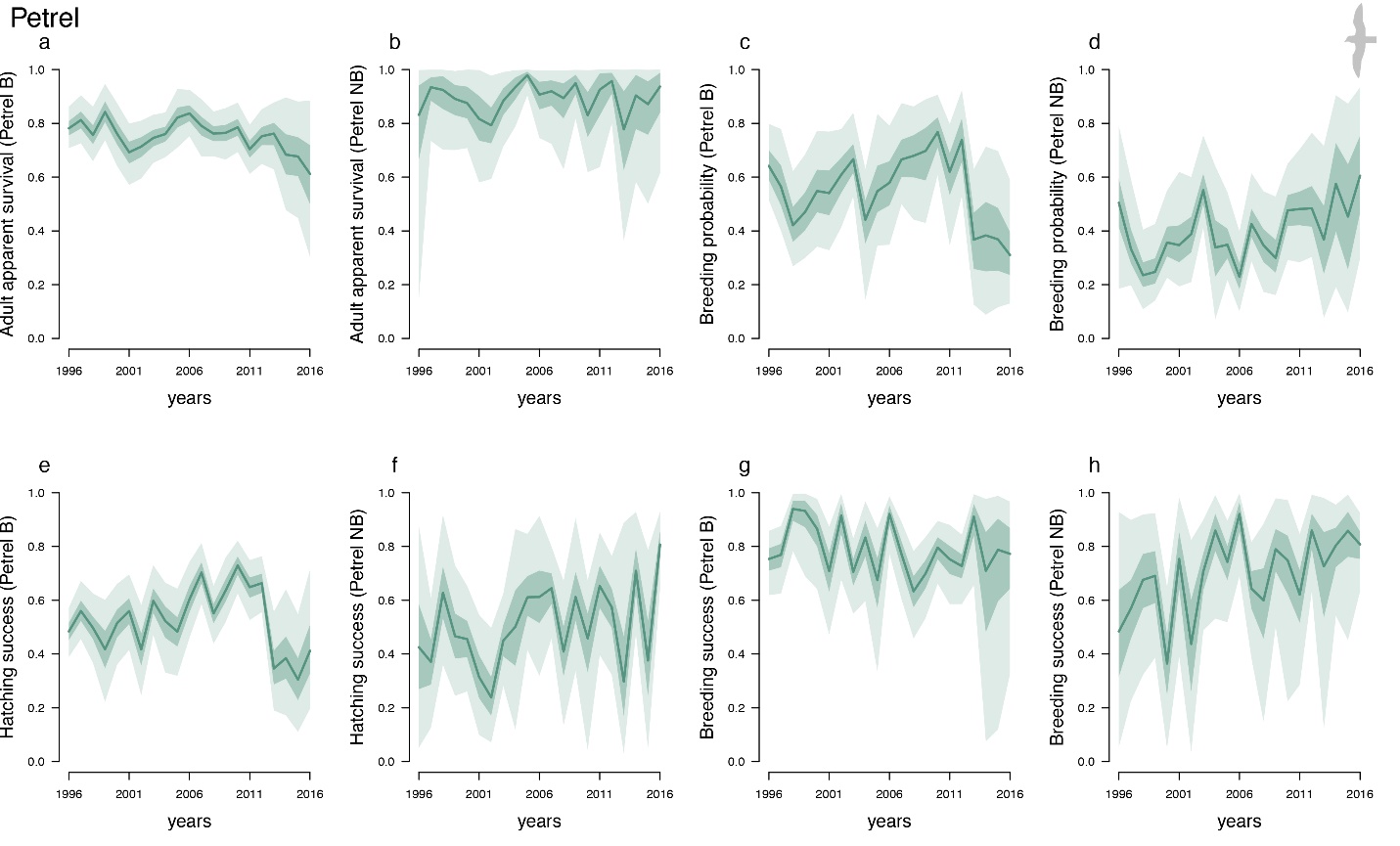
Figure S2: Demographic parameter estimates for each year (1996-2016) for Blue Petrels. Solid line represent the means of marginal posterior distributions. Shaded area are the 50% and 95% credibility intervals. (a, b) Estimated adult apparent survival probability$\phi_{P,\tilde{B}}$ and $\phi_{P,\tilde{NB}}$ (c, d) Estimated breeding probability$\beta_{P,\tilde{B}}$ and $\beta_{P,\tilde{NB}}$ (e, f) Estimated hatching success probability $\omega_{P,\tilde{B}}$and $\omega_{P,\tilde{NB}}$ (g, h) Estimated breeding success probability $\gamma_{P,\tilde{B}}$ and $\gamma_{P,\tilde{NB}}$ for petrel that have bred ($\tilde{B}$) or have not bred ($\tilde{NB}$), respectively, during the previous breeding season.

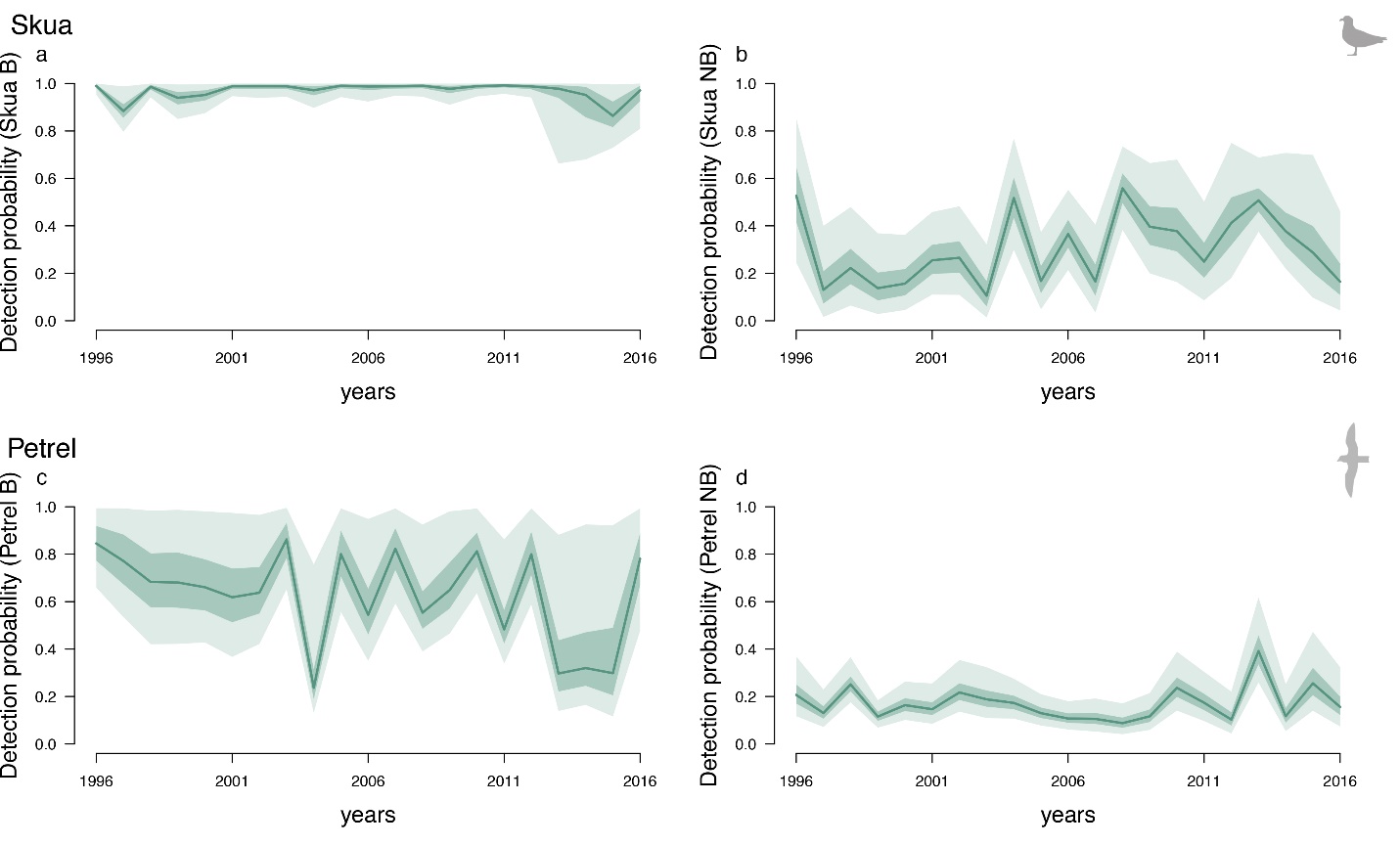

Figure S3: Detection probability estimates for each year (1996-2016) for Brown Skuas (top) and Blue Petrels (bottom). Solid lines represent the means of marginal posterior distributions. Shaded area are the 50% and 95% credibility intervals. Estimated detection probability for (a) breeder ${(p}_{S,\tilde{B}})$and (b) nonbreeder ${(p}_{S,\tilde{NB}})$ skuas, (c) breeder ${(p}_{P,\tilde{B}})$and (d) nonbreeder $(p_{P,\tilde{NB}})$petrels.

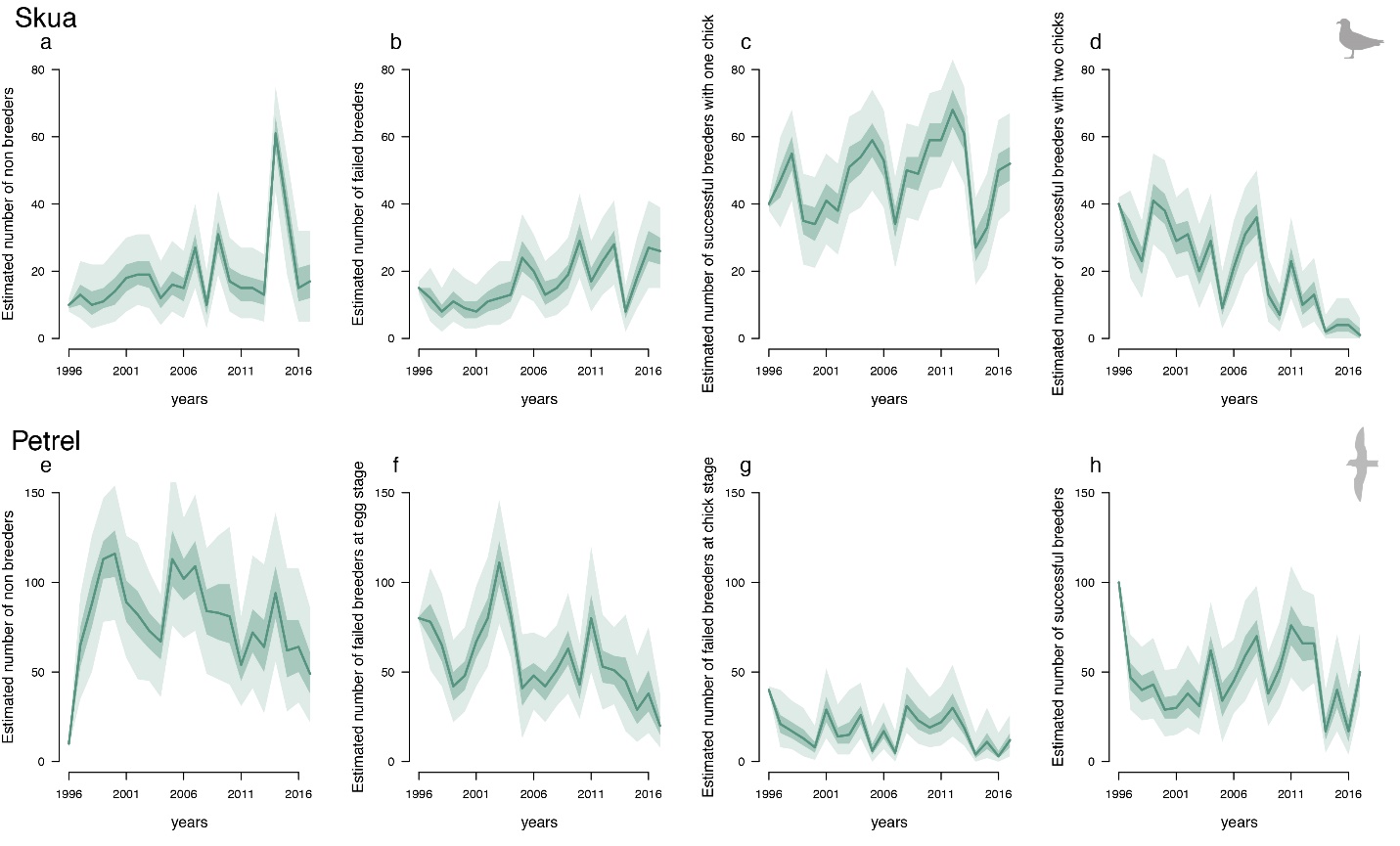

Figure S4: Estimated number of individuals in adult states for each year (1996-2016) for Brown Skuas (top) and Blue Petrels (bottom). Solid lines represent the means of marginal posterior distributions. Shaded area are the 50% and 95% credibility intervals. Estimated number of skuas (a) nonbreeders $N_{NB,S}$ (b) failed breeders $N_{FB,S}$ (c) successful breeders with at least one chick $N_{SB1,S}$ (d) successful breeders with two chicks$N_{SB2,S}$. Estimated number of petrels (e) nonbreeders $N_{NB,P}$ (f) failed breeders at egg stage $N_{FBE,P}$ (g) failed breeders at chick stage $N_{FBC,P}$ (h) successful breeders$N_{SB,P}$.

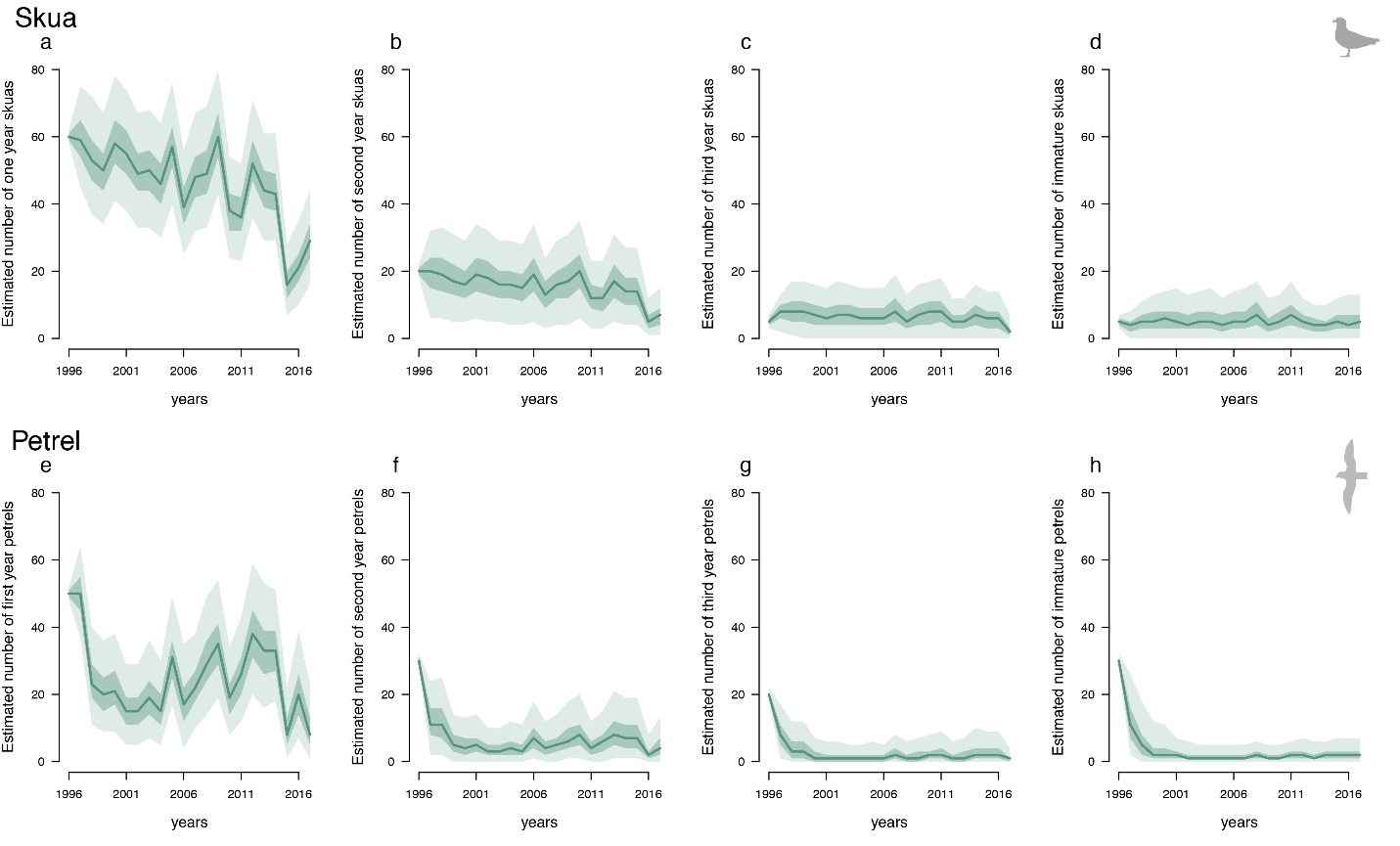

Figure S5: Estimated number of individuals in juvenile states for each year (1996-2016) for Brown Skuas (top) and Blue Petrels (bottom). Solid lines represent the means of marginal posterior distributions. Shaded area are the 50% and 95% credibility intervals. Estimated number individuals (a, e) in their first year $N_{J1,S}$ and $N_{J1,P}$ (b, f) in their second year $N_{J2,S}$ and$N_{J2,P}$ (c, g) in their third year$N_{J3,S}$ and$N_{J3,P}$, (d, h) in their fourth year or more (immatures) $N_{J4+,S}$ and $N_{J4+,P}$ for skuas and petrels, respectively.

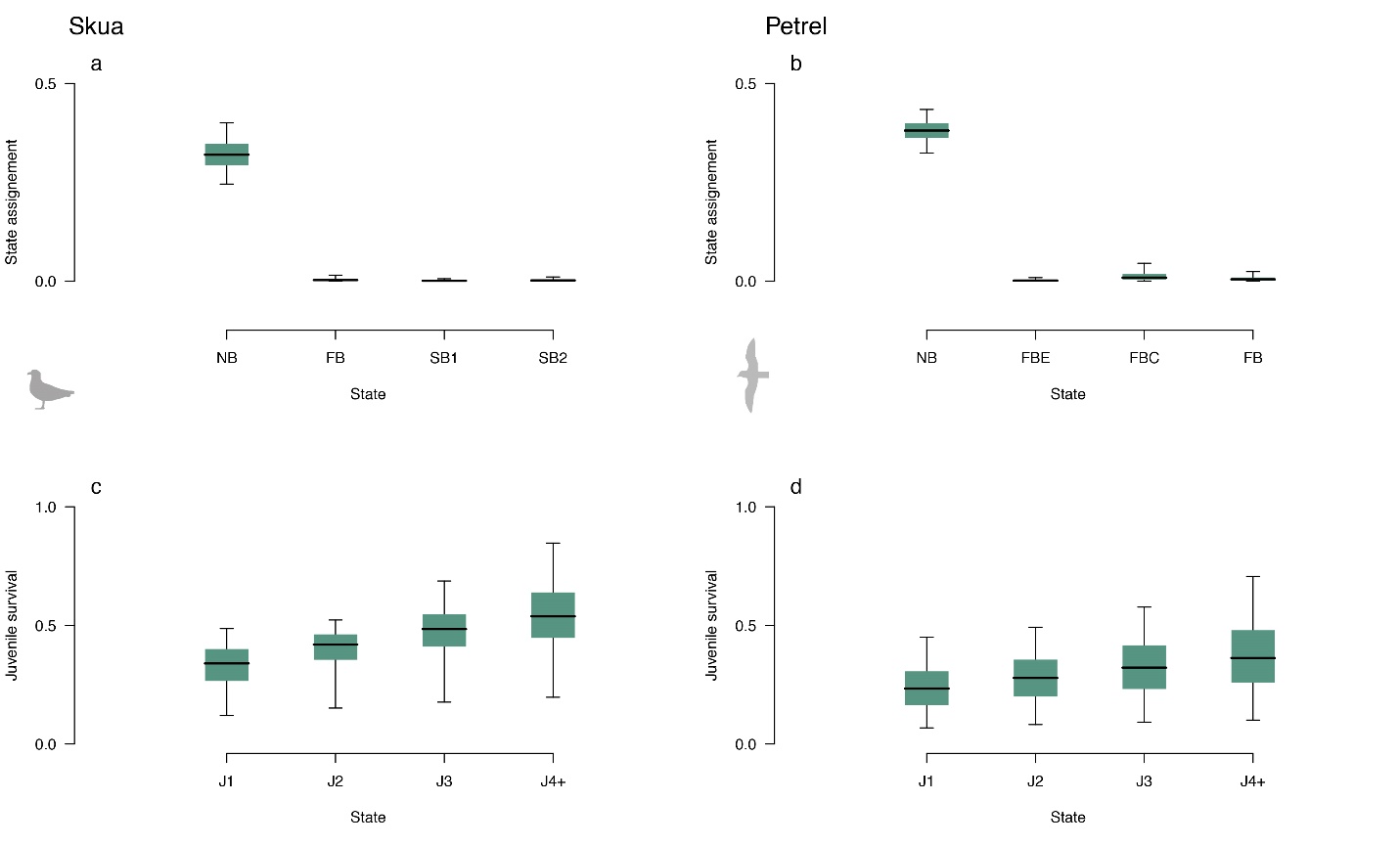

Figure S6: Estimates of (a, b) state assignment probability $(1-u)$ for Nonbreeder (NB), Failed breeder (FB) at the egg stage (FBE), at the chick stage (FBC), the successful breeders (SB) with one (SB1) or two (SB2) chicks. (c, d) juvenile apparent survival probability (time independent) $\phi_{J}$ for juvenile birds of one year old ($J1$), two years old ($J2$), three year old ($J3$), and four years old and older ($J4+$) for Brown Skuas (left) and Blue Petrels (right). The black line corresponds to the mean of marginal posterior distributions, the color interval corresponds to the 50% credibility interval and the arrows correspond to the 95% credibility interval.
